# Neural mechanisms of training in Brain-Computer Interface : A Biophysical modeling approach

**DOI:** 10.1101/2025.06.21.660834

**Authors:** Apurba Debnath, Tristan Venot, Marie-Constance Corsi, Parul Verma

## Abstract

Brain-Computer Interface (BCI) is a system that translates neural activity into commands, allowing direct communication between brain and external devices. Despite its clinical application, BCIs are unable to robustly capture subjects’ intent due to limited understanding of the neural mechanisms underlying BCI control. To address this, we introduce a biophysical linear neural mass model to investigate the associated neural mechanisms of motor imagery BCI (MI-BCI) by modulating the scale-free spectral characteristics of afferent inputs. We tailor this motor imagery neural mass model (mi-NMM) to simulate both MI and resting-state activity and apply it to a cohort of 19 healthy subjects trained over four sessions, during which magnetoencephalography and electroencephalography signals were simultaneously recorded. Task-dependent significant modulation of scale-free cortical dynamics reveals training-induced excitatory–inhibitory (E/I) reconfiguration during BCI training, shifting from widespread cortical to targeted sensorimotor recruitment. The modeled intra-regional E/I interaction strengths and excitatory population time constants capture condition- and session-related changes in neural activation in the relevant areas of the sensorimotor cortex, where MI-related desynchronization and synchronization are associated with excitatory population time constants. Thus, these mechanistic findings provide new insights into the underlying neural mechanisms of MI-BCI, paving the way for tailored BCI training protocols.

## Introduction

Brain-Computer Interfaces (BCIs) enable the brain to directly control external devices by translating neural activity into actionable commands, but developing a control system based on brain signals is a formidable challenge. BCIs necessitate the acquisition of intricate neural data and the conversion of these signals into dependable commands, all within the constraints of an incomplete understanding of brain function (1; 2). Despite these obstacles, BCIs hold significant promise, particularly in clinical applications, where they enable direct brain-to-device communication to compensate for lost motor or sensory functions (3).

Nonetheless, non-invasive methods such as electroencephalography (EEG) and magnetoencephalography (MEG) often provide limited control for many users (4). While training can enhance BCI performance, a critical question persists: why do some users exhibit substantial improvement while others face ongoing difficulties? This issue is particularly evident in motor imagery-based BCIs (MI-BCI), where users operate a device by imagining movements without physically executing them. Numerous studies have attempted to address the various ambiguities in this field, including enhancing brain signal decoders (5), identifying neurophysiological and psychological factors affecting user performance (6; 7; 8), and developing improved training protocols (9). A critical element believed to be essential for successful BCI control is neuroplasticity, which underlies the brain’s capacity to adapt and learn. This has spurred increasing interest in the neurophysiological mechanisms involved in neurofeedback and BCI learning (10). Furthermore, the phenomenon of BCI inefficiency has been examined from multiple perspectives. Learning to use a BCI differs fundamentally from interacting with intuitive, everyday tools; it requires mastering a new, abstract form of control. It is reasonable to infer that specific brain mechanisms are involved when a user learns to perform BCI task effectively.

Learning is increasingly understood not merely as a localized change in neural activity, but as a reconfiguration of the brain’s overall dynamic state (11). A key signature of this state is the non-oscillatory brain activity, termed as scale-free/aperiodic activity for the lack of definitive temporal scale, reflected in the electrophysiological power-law relationship of 1*/f*^*k*^ across frequencies, quantified by the spectral exponents value (*k*) and it is thought to arise from coordinated synaptic inputs. Importantly, this scale-free/aperiodic activity is not static; rather, it changes dynamically with task demands, cognitive states and pathological alterations in neurological disorders (12; 13; 14). However, it remains unclear how the task-driven shifts in aperiodic activity contribute to the reorganization of excitatory-inhibitory (E/I) interactions during learning, underscoring need for biophysical models capable of elucidating the E/I dynamics involved in BCI learning and adaptation, while accounting for shifts in aperiodic activity.

A class of biophysical models that have been particularly useful for these purposes is called the “Neural Mass Models” (NMMs) (15). NMMs model the population-level activity of interactions between excitatory and inhibitory neurons in different brain regions that potentially underlies the macroscopic signals we observe with M/EEG. As a result, they provide biophysical insights into how macroscopic signals may arise in various neurological conditions/diseases. Although these models have been used widely, they have rarely been investigated in the case of BCI, potentially because of the additional challenge of modeling the tasks appropriately. Indeed, most modeling studies focus on the resting state (RS) with limited attempts to model BCI-related tasks (16; 17). In this study, we developed a novel approach to model MI-BCI, called mi-NMM, that explicitly accounts for the brain’s scale-free dynamics, capturing the prevalent arrhythmic activity that co-exists, yet is distinct from, brain oscillations. It leverages a linearized NMM that accurately captures the spectral features of the neural oscillatory activity estimated from source-localized MEG in healthy individuals in the RS (18; 19). This NMM has also been successful in identifying cortical-level neural impairments in patients with Alzheimer’s disease (20) as well as neural modulations in healthy individuals transitioning from wakefulness to light sleep (21).

Using this approach, we hypothesized that key spectral differences between MI and RS result from region-specific, population-level interactions between excitatory and inhibitory neural populations. Furthermore, we posited that such region-specific changes could capture the neural mechanisms occurring during learning. To validate these hypotheses, we utilized source-reconstructed EEG and MEG signals obtained during a four-session MI-BCI training from healthy subjects. A schematic of our approach is being shown in Fig.1.

**Fig. 1.**
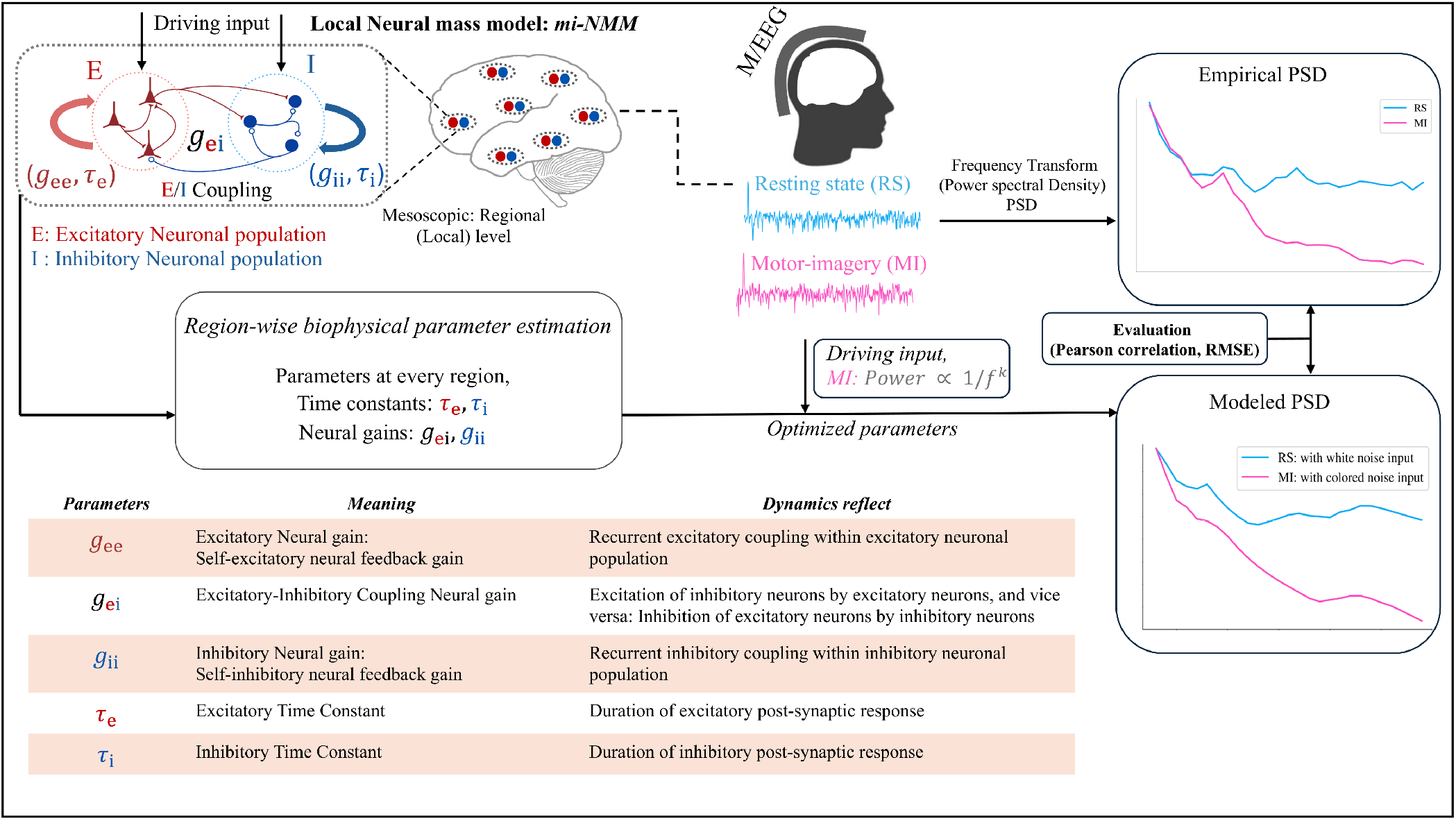
Schematic of the modeling approach. On the top left, we show that the excitatory and inhibitory neuronal populations are modeled for every brain region using a small set of biophysically meaningful parameters (summary table at the bottom left). Both these populations also receive a driving input that is simulated as white noise, characterized by frequency-independent power for resting state (RS), and as colored noise, characterized by frequency-dependent power law form of (1*/f* ^*k*^), during motor imagery (MI). Upon simulating the model, we directly obtain the modeled PSD, shown at the bottom-right and compare with the empirical PSD at top-right and optimize the model parameters (shown in the middle) by matching the modeled PSD to the M/EEG PSD for every region. After obtaining the optimized parameters, we performed statistical analyses to test for condition and session effect in both M/EEG separately, map the parameters distribution across brain, and relate those parameters to the training-induced changes in E/I dynamics.

## Results

### Motor-imagery condition is accompanied by significant increase in aperiodic spectral exponents

First, to compare the aperiodic activity between the MI and RS conditions, we calculated the spectral exponents from the empirical spectra using FOOOF, a Python package (22) designed to fit to the power spectra from M/EEG recordings and calculate the aperiodic spectral exponent values. The results showed that FOOOF provided a better fit to the median spectra across subjects (calculated by taking the median across all subjects for each ROI) than to individual spectra in almost all the sessions/modalities that we considered, based on the R2 metric measuring the goodness-of-fit between the actual spectra and the FOOOF-modeled spectra. For MEG, the mean R2 values for RS condition were 0.983 (median spectra) vs. 0.950 (individual spectra), and for MI condition were 0.990 (median spectra) vs. 0.980 (individual spectra). For EEG, the mean R2 values for RS condition were 0.859 (median spectra) vs. 0.865 (individual spectra), and for MI condition were 0.973 (median spectra) vs. 0.9118 (individual spectra). This was likely because the median spectrum was smoother, less noisy and therefore, easier to fit to using FOOOF, rather than some of the individual spectra. Note that we used the median instead of the mean when comparing the spectra, as there were few potential outliers that could skew the results (see Fig. S1). Our results revealed that the MI was accompanied by significant increase in aperiodic spectral exponents compared to RS condition, in all sessions and imaging modalities considered in this study, with a very strong effect size (MEG: Paired-t Test: *t* = −5.36, *p* = 1.1 × 10^*−*6^, Cohen’s *d*_*z*_ = 0.645; EEG: Paired-t Test: *t* = −6.18, *p* = 4.18 × 10^*−*8^, Cohen’s *d*_*z*_ = 0.75) and clearly shown in Fig. 2.

**Fig. 2.**
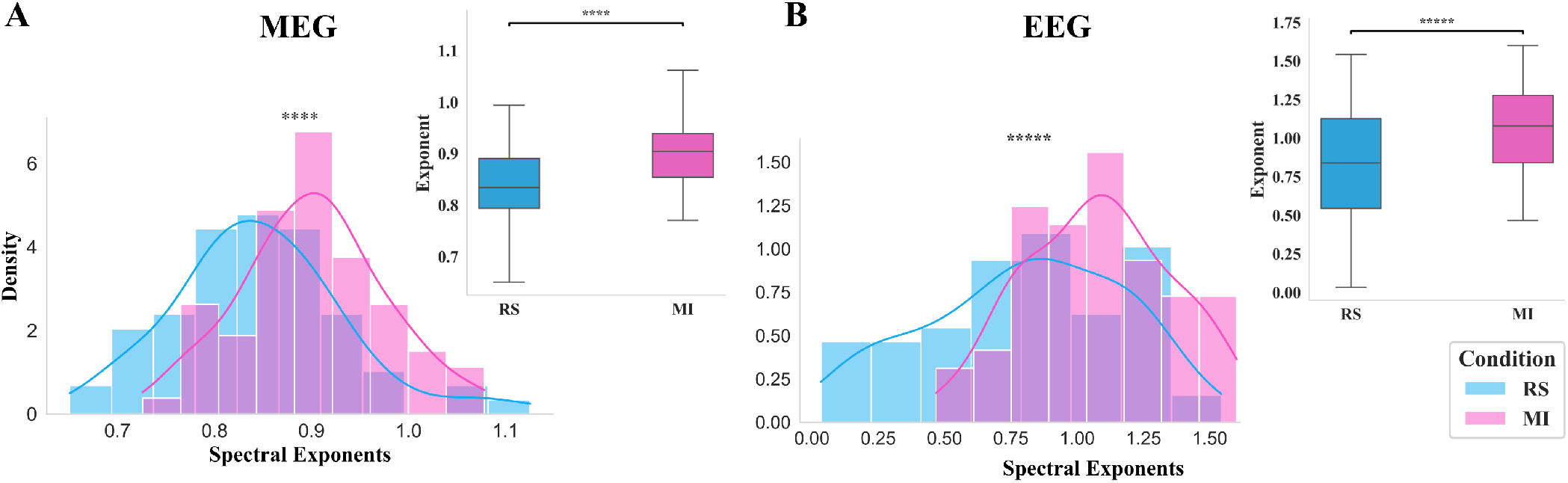
Motor-imagery (MI) is characterized by a significant increase in aperiodic spectral exponents. (A, B) exhibit group-level median spectral exponents in both MEG, and EEG modalities, respectively, and demonstrate that the aperiodic spectral exponents during MI are significantly increased compared to during RS. Compared to RS, MI exhibits a significant shift toward larger spectral exponents in both modalities, with a very strong effect size (MEG: Paired-t Test: *t* = −5.36,****: *p* = 1.1 × 10^*−*6^, Cohen’s *d*_*z*_ = 0.645; EEG: Paired-t Test: *t* = −6.18,*****: *p* = 4.18 × 10^*−*8^, Cohen’s *d*_*z*_ = 0.75), indicating task-dependent significant modulation of scale-free cortical dynamics.

### Simulating modeled input as colored noise better captures MI power spectra in M/EEG

The NMM for the excitatory/inhibitory neural population signals incorporates decay of these signals over time, interactions among them, and a driving input *p*(*t*). In our framework, this driving input/stimulus refers to aggregate latent background drive received by each neural population, which encompasses influences that are not explicitly modeled within the local neural mass, including both intrinsic background activity and extrinsic afferent inputs arising from distributed cortical/subcortical regions through the structural connectome. Under RS condition, this aggregate drive is commonly approximated to be largely asynchronous and temporally uncorrelated stochastic input. Accordingly, following previous studies (18; 20; 19; 23; 24; 25; 16), we modeled this input *p*(*t*) as white noise for RS condition in Equation (6), characterized by frequency-independent power spectral density (PSD), for all ROIs, subjects, and sessions.

Our empirical analysis (Section 3.1) demonstrated that MI was accompanied by a significant increase in aperiodic spectral exponents compared to RS, with a strong effect size, indicating stronger scale-free temporal correlations in the measured macroscopic M/EEG signals during MI condition. Thus, we used it as a parsimonious approximation of the temporally correlated latent background drive contributing to the macroscopic M/EEG signal during MI. To this end, we modified spectral characteristics of this input when simulating the MI condition. Specifically, *p*(*t*) was modeled as colored noise, characterized by aperiodic power-law form of 1*/*(*f* ^*k*^), where the exponent *k* was allowed to vary within the range [0, 1] for all ROIs, subjects, and sessions (*ω* = 2*πf*). We call this modified model as *mi-NMM*.

Moreover, this modeling approach accurately reproduced the empirical MI power spectra, with more than 90% of all fitted modeled spectra exhibiting an RMSE below 0.3. In comparison, the RS model produced RMSE values extending up to 0.5 over an equivalent number of spectral evaluations as shown in Fig. 3-(E, F) with both M/EEG imaging modalities.

**Fig. 3.**
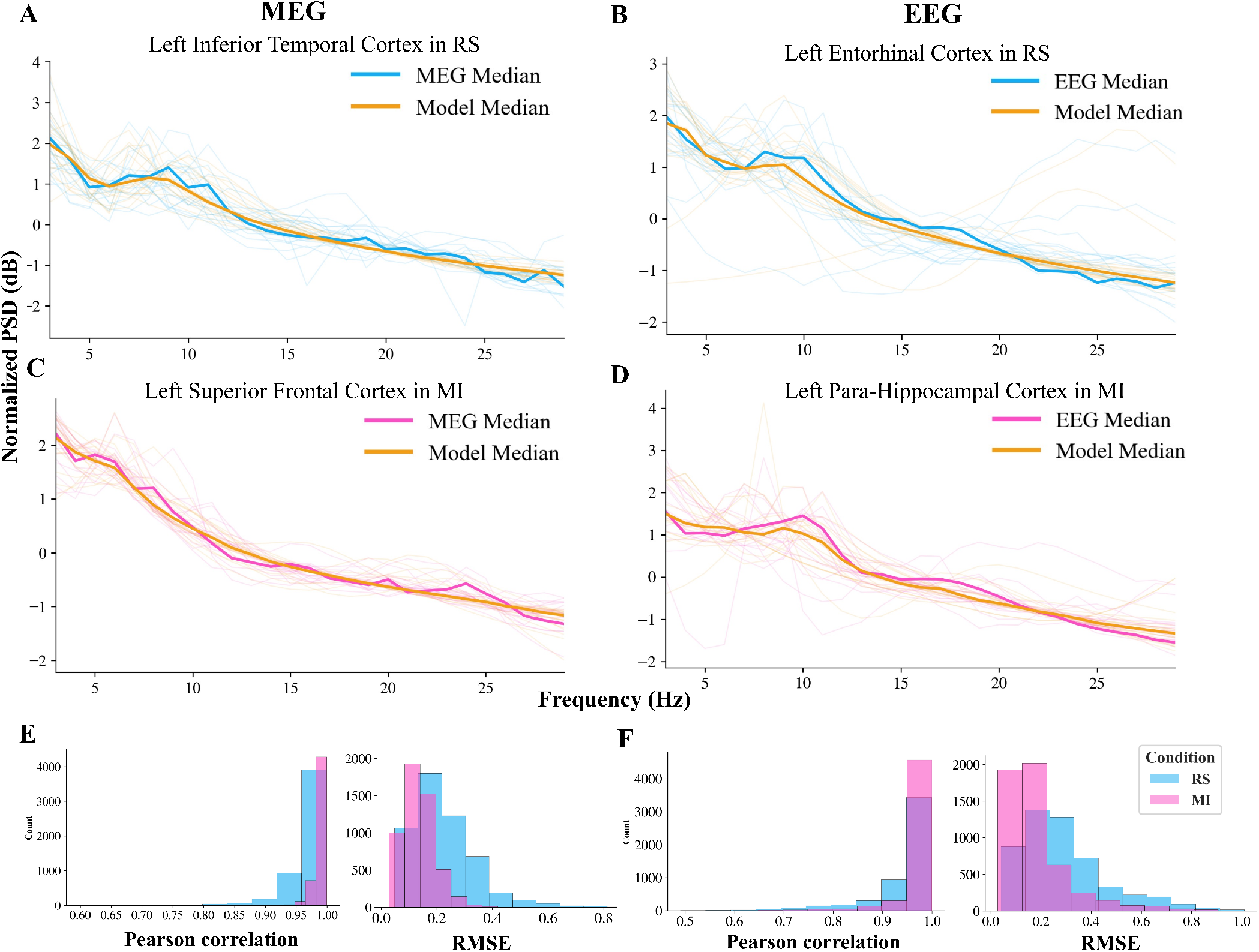
mi-NMM mirrors M/EEG power spectra accurately in both MI and RS conditions. Plot (A, B) show the empirical and modeled PSDs with high Pearson correlations ∈ [0.85, 0.99] and RMSE ∈ [0.1, 0.5] for the Left Inferior Temporal cortex and Left Entorhinal cortex in all 19 subjects in RS condition with MEG and EEG imaging modalities, respectively. Bold spectra represent modeled median and empirical median PSDs across subjects. Similarly, (C, D) show the empirical and modeled PSDs with high Pearson correlations ∈ [0.85, 0.99] and RMSE ∈ [0.07, 0.2] for the Left Superior Frontal cortex and Left-Hippocampal cortex in MI condition, respectively. The overlap between these median curves clearly highlights how well the model captures the shape of the actual spectra. (E, F) show the distribution of Pearson correlation coefficients and RMSE between the modeled and actual PSDs for all ROIs, subjects, and sessions we used in our study, containing both MI and RS conditions for MEG and EEG, respectively. In most cases, Pearson correlation exceeds 0.90, with 90% of the spectra showing RMSE lower than 0.3 during MI, while a slightly higher RMSE is observed in RS condition with MEG and EEG.

### The biophysical mi-NMM captures M/EEG power spectra accurately in both RS and MI conditions

The comparison between the modeled and actual PSD (Z-score Normalized) across different brain regions in all subjects and sessions revealed a strong correspondence in the decibel (dB) scale over the frequency range of 3 to 30 Hz. The spectral shape of the modeled PSD closely aligned with that of the actual PSD across all examined regions, subjects, and sessions, demonstrating the effectiveness of the modeling approach in capturing neural oscillatory patterns measured with M/EEG, while accounting for shift in scale-free/aperiodic brain activity during MI. To demonstrate this qualitatively, example spectra for a subset of regions in all subjects are shown in Fig. 3 (A-D). Here, the regions for RS-modeled MEG (A, Left inferior temporal cortex), RS-modeled EEG (B, Left entorhinal cortex), MI-modeled MEG (C, Left superior frontal cortex), and MI-modeled EEG (D, Left para-hippocampal cortex) are shown, respectively, with high accuracy (Pearson correlation ∈ [0.85, 0.99], RMSE ∈ [0.1, 0.5]). Moreover, quantitative assessment done by two goodness-of-fit metrics: Pearson correlation and RMSE on the modeled spectra for all 68 cortical regions, subjects, and sessions reinforced these findings. The Pearson correlation coefficient between the modeled and actual PSD exceeded 0.90 in most cases in both RS and MI for M/EEG. The RMSE values were also below 0.2 in most cases in MI, with slightly higher values in the RS condition (Fig. 3 - E, F). This high correlation indicates that mi-NMM derived from biophysically realistic parameters’ range successfully replicated key spectral features captured via M/EEG in different sessions and regions.

### Sensitivity analysis reveals time constants strongly influence modeled spectra while input exponent shape spectral slope

Thereafter, to analyze the influence of the model parameters and input exponent on modeled spectra generation, we performed local sensitivity analysis across all ROIs, subjects, and sessions for both RS and MI conditions, in both MEG and EEG. Sensitivity was quantified by the spectral sensitivity coefficient for each model parameter and input exponent. A larger sensitivity coefficient indicates a greater influence of a parameter/exponent on the generated modeled spectral profile. We repeated the analysis for several perturbation values (*ϵ*) and in all analyses we found that the time constants (*τ*_e_, *τ*_i_) had the most influence on model spectra generation, while neural gains (*g*_ii_, *g*_ei_) had less influence. The excitatory time constant (*τ*_e_) exhibited the highest sensitivity in both RS and MI, indicating that the modeled spectra were most sensitive to perturbations in *τ*_e_. An example perturbation of 10%(*ϵ* = 0.10) in the model parameters and input exponent showed that *τ*_e_ produced the largest relative spectral changes during both RS and MI, with the mean spectral sensitivity coefficient of 1.58 (MEG) and 1.81 (EEG) during RS (Fig. 4-A, B); 1.29 (MEG) and 1.59 (EEG) during MI (Fig. 4-C, D). In contrast, the smallest changes during RS were observed for *g*_ii_ (0.34 in MEG) and *g*_ei_ (0.36 in EEG), whereas during MI, the input exponent *k* showed the lowest sensitivity, with sensitivity coefficients of 0.16 in MEG and 0.27 in EEG. For additional information, see Table S1 & S2.

**Fig. 4.**
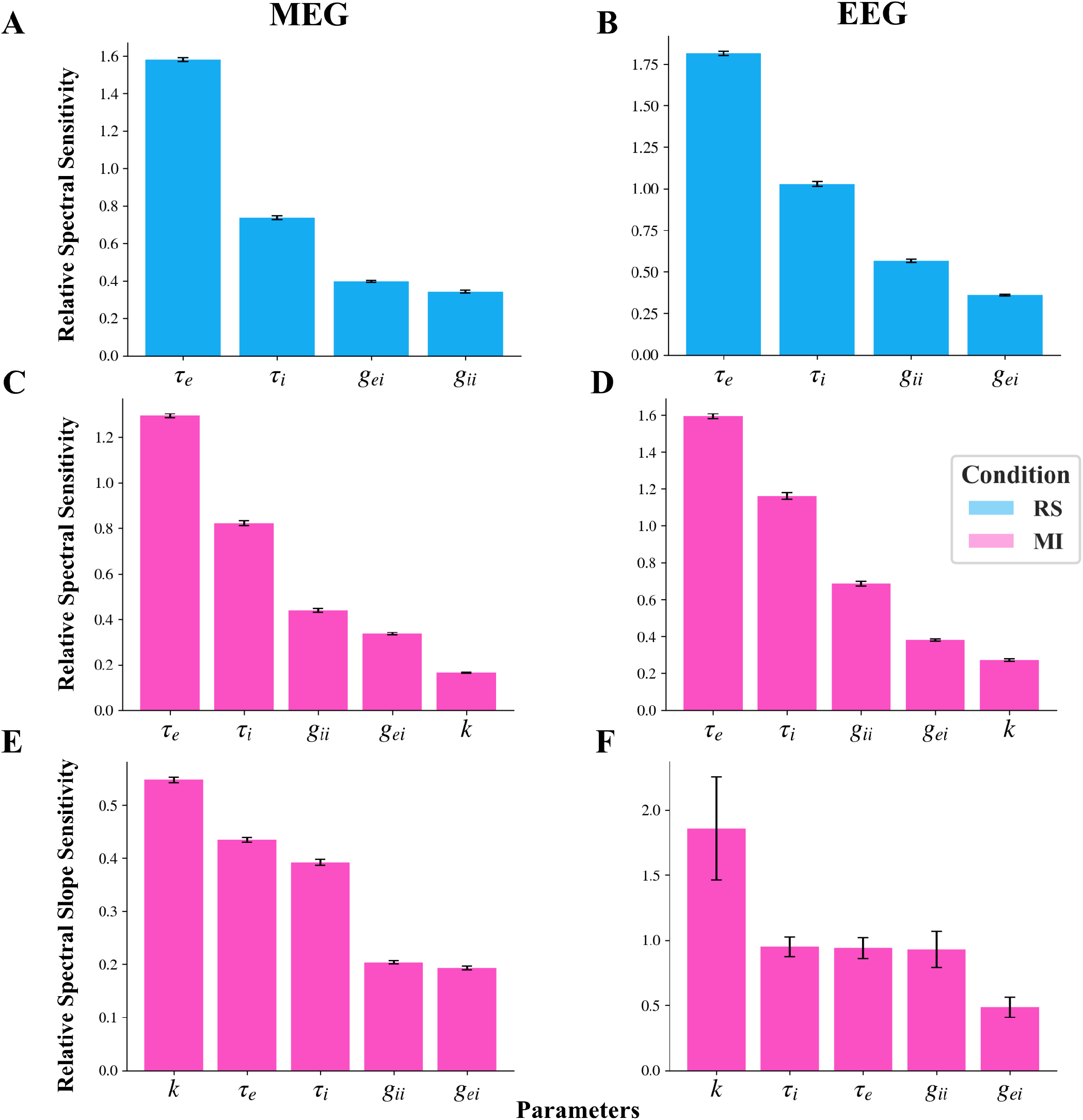
Influence of model parameters and input exponent on the generated modeled PSD and its slope. (A, B) show relative spectral sensitivity coefficients of the model parameters and input exponent for the modeled MEG and EEG PSD during RS, respectively, computed across all ROIs, subjects, and sessions. Similarly, (C, D) show sensitivity coefficients during MI with MEG and EEG, respectively. The excitatory time constant (*τ*_e_) shows the highest spectral sensitivity toward modeled spectra during both RS and MI, with MEG/EEG, indicating that the modeled spectra are most sensitive to perturbations in *τ*_e_. Here, perturbation by 10%(*ϵ* = 0.10) in model parameters and input exponent show that *τ*_e_ produces most relative spectral changes with mean coefficient of 1.58 (A-MEG), 1.81 (B-EEG) during RS and 1.29 (C-MEG), 1.59 (D-EEG) during MI. In contrast, the smallest changes are observed for neural gains: *g*_ii_ (0.34, A-MEG) and *g*_ei_ (0.36, B-EEG) during RS, compared to during MI where the input exponent shows least sensitivity (*k*: 0.16, C-MEG; 0.27, D-EEG) towards the modeled PSD generations. (E, F) show the sensitivity of model parameters and input exponents towards modeled spectral slope during MI in MEG and EEG, respectively. The input exponent (*k*) has the greatest influence on slope compared to model parameters. An example perturbation by 20%(*ϵ* = 0.20) shows that *k* has the most influence on slope with the mean slope sensitivity of 0.54 in MEG (E) and 1.85 in EEG (F) compared to model parameters.

Additionally, our slope sensitivity analysis revealed that the input exponent (*k*) had the maximum influence on the modeled spectral slope. An example perturbation by 20%(*ϵ* = 0.20) showed that the input exponent (*k*) exhibited the highest mean spectral slope sensitivity (MEG, EEG) with (0.54, 1.85), followed by model parameters: *τ*_i_ (0.39, 0.95), *τ*_e_ (0.43, 0.94), *g*_ii_ (0.20, 0.93), and *g*_ei_ (0.19, 0.49), indicating that the input exponent had the most influence on the modeled spectral slope during MI, shown in Fig.4-E, F. This was the case for all the *ϵ* values used in this study. For further details, refer to Table S3 & S4.

### BCI training drives a shift from diffuse to targeted sensorimotor activation via regional E/I dynamics

The neurophysiologically meaningful model parameters reflect the underlying E/I population-level dynamics that shape how and when a region becomes active in response to intrinsic or extrinsic driving inputs under different conditions. Thus, to investigate the possible condition- and session-related modulation/effect for MI versus RS, we conducted the Wilcoxon signed-rank test for each of these parameters and regions separately in each session and applied FDR correction for multiple comparisons, and found significant parameter-related modulation (*p*_FDR_ *<* 0.05) for different brain regions with very strong effect sizes (*r* ∈ [0.6, 0.8]). We first examined the spatial distribution of significant model parameters in MEG: neural gains (*g*_ei_, *g*_ii_), and time constants (*τ*_e_, *τ*_i_) with a focus on a shift in activation patterns grouped by lobe. We specifically did not find any significant region for *τ*_i_ (see Fig. 5). For an exhaustive presentation of the regions that showed significant effects, please refer to the supplementary table (Table S5) with the list of significant regions of interest (ROIs), session by session, for each of these parameters. Below, we briefly describe these findings in motor and non-motor regions separately.

**Fig. 5.**
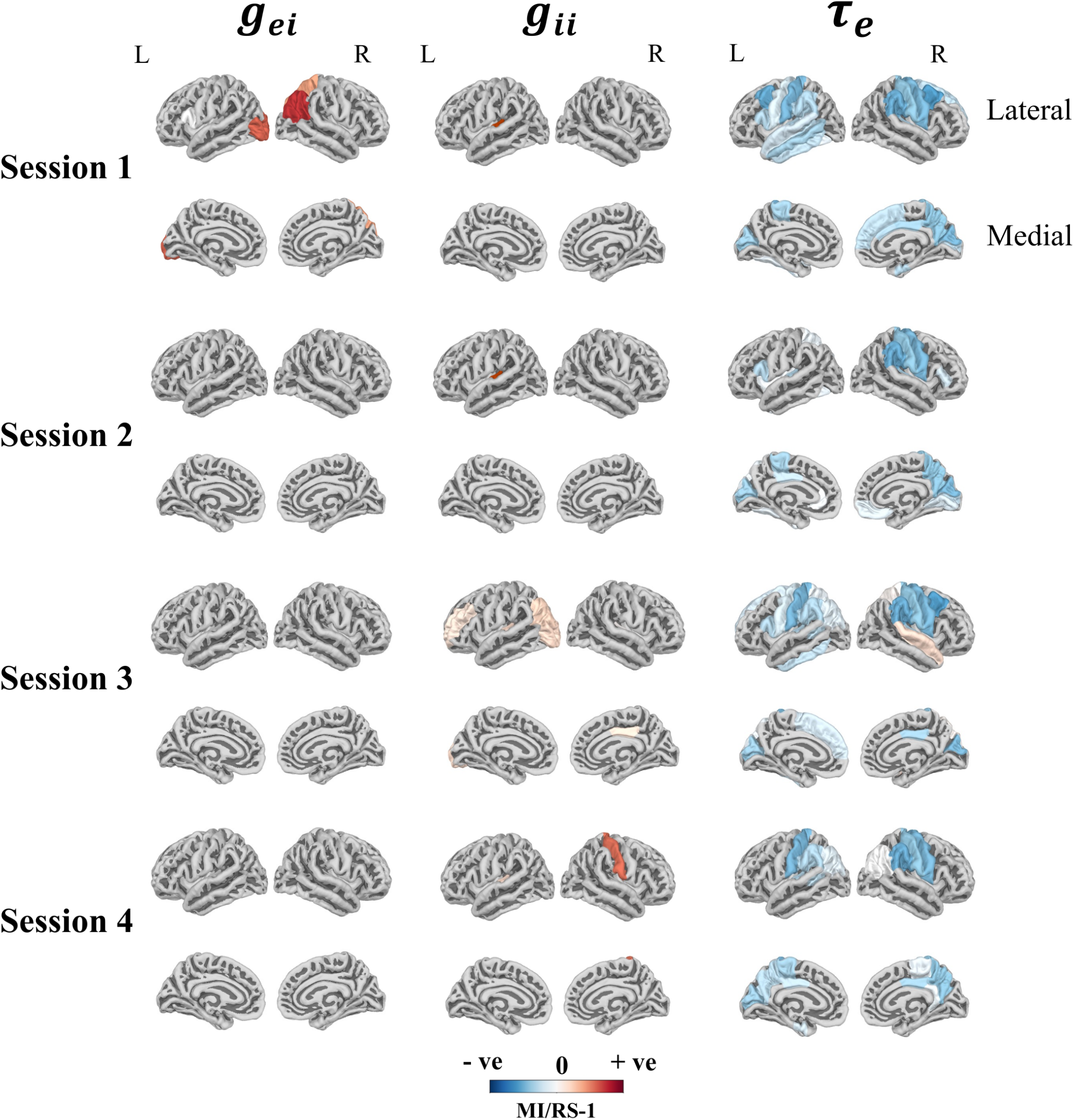
BCI training drives a shift from diffuse to targeted sensorimotor activation – refinement of cortical recruitment. The colored regions, described by mean (MI/RS-1) across subjects, show significant condition-related modulation with a very strong effect size (*p*_FDR_ *<* 0.05; Effect size, *r* ∈ [0.6, 0.8]) for MI Vs RS, with the three MEG fitted model parameters (*g*_ei_, *g*_ii_, *τ*_e_), respectively, across the four training sessions. Early training sessions exhibit widespread and distributed cortical involvement, whereas later sessions show more selective engagement of task-relevant sensorimotor regions, robustly captured by excitatory time constant *τ*_e_. This progressive reduction in spatial extent from 29 regions in session-1 to 19 regions in session-4 reflects a shift from diffuse to targeted cortical recruitment with training. The minimum and maximum range of mean (MI/RS-1) for the parameters across sessions are (−0.42, +158.47), (+1.95, +336.08), (−0.354, +0.628) for (*g*_ei_, *g*_ii_, *τ*_e_) respectively.

As subjects were instructed to perform motor imagery tasks, our primary interest was to investigate the potential involvement of motor areas. For *g*_ii_, which reflects self-inhibitory neural feedback gain within the inhibitory population, the *right postcentral gyrus* was found to be accompanied by significantly higher *g*_ii_ during MI by session 4 (Fig. 5), reflecting disinhibition. The parameter *τ*_e_, which characterizes the duration of the excitatory post-synaptic response, yielded the most extensive and consistent activation patterns across sessions with the dynamic recruitment of the *paracentral*, the *precentral*, and the *postcentral* cortex during the training, accompanied by significant reduction of *τ*_e_ values in MI compared to RS in different sessions (Fig. 5), including the left motor areas (Fig. 6, Wilcoxon signed-rank test: Bonferroni corrected, *p <* 0.05; Effect size, *r* ∈ [0.6, 0.8]). Additionally, the left *paracentral* cortex, which showed a strong modulation of *τ*_e_ in session-4, also exhibited a weak training effect for *τ*_e_ without correction for multiple testing (Friedmann test: uncorrected *p <* 0.05, Bonferroni corrected *p* = 0.0904). More specifically, such an effect was driven by the MI condition and not RS condition (see Fig. 6-A). This change suggests that neural activity in the left paracentral region may be dynamically influenced between sessions during MI, although the effect size was weak due to limited training sessions (Kendall’s W = 0.14). Moreover, our LMM analysis for all 68 regions, while considering subject as random intercept, revealed a modest positive correlation between *τ*_e_ during MI and group-level BCI performance across sessions for the left precentral cortex (Fig. 6-E, LMM: *β* = 0.38, uncorrected *p <* 0.05). Even though the *τ*_e_ was weakly correlated with group-level BCI performance during MI, and this was not significant after multiple comparisons corrections, we still report this finding here, as it raises the possibility that *τ*_e_ may capture the sustained involvement of motor regions later during training, given that it was already significantly modulated in multiple motor regions by Session 4, as shown in Fig. 6.

**Fig. 6.**
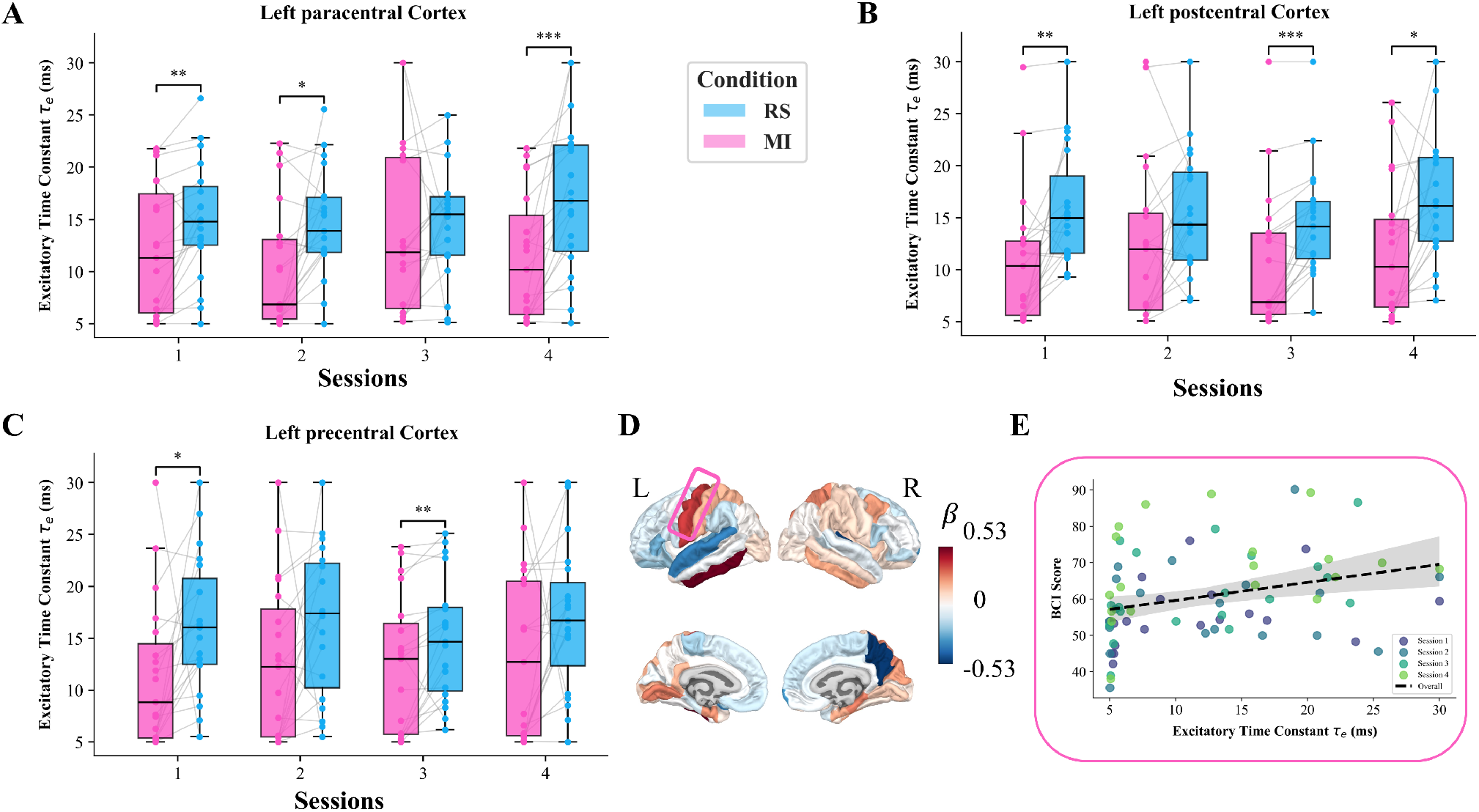
Dynamic complementary activation of motor cortex areas across sessions is reflected by the excitatory time constant (*τ*_e_). (A, B, C) show the distribution of 19 individual subjects’ *τ*_e_ values during MI and RS for each session in left paracentral, postcentral, and precentral cortex areas, respectively. All 3 regions are activated in session-1, accompanied by a significant reduction in *τ*_e_ values for most subjects during MI (Wilcoxon signed-rank test: Bonferroni corrected, *: *p <* 0.05,**: *p <* 0.01; Effect size, *r* ∈ [0, 6, 0.7]). Session-2 shows activation of left paracentral area, whereas in session-3, activation shifts toward the left postcentral and precentral areas, and then in the last session-4, left paracentral area is strongly activated by a significant reduction in *τ*_e_ values (Bonferroni corrected, ***: *p <* 0.001; Effect size, *r* = 0.8). Additionally, A shows inter-session variation in the left paracentral region driven by MI condition (Friedamann test: uncorrected *p <* 0.05, Bonferroni corrected *p* = 0.0904), and not RS condition. Plot D, spatial map shows the *β* coefficient from the LMM analysis on all regions, relating the *τ*_e_ to group-level BCI scores across sessions during MI condition, where the positive values (warm colors) indicate that higher *τ*_e_ is associated with higher BCI scores, whereas negative values (cool colors) indicate the opposite during MI. Plot E shows modest positive correlation across sessions between the *τ*_e_ and group-level BCI performance for the left precentral cortex driven by MI condition (LMM: *β* = 0.38, uncorrected *p <* 0.05).

We observed the involvement of non-motor regions throughout the training, beyond those targeted by the BCI system (Fig. 5). With *g*_ei_ parameter that captures recurrent interactions among E/I neurons, effects were exclusively observed in session 1, with regions in the frontal lobe (left *pars opercularis*), the parietal lobe (right *inferior* and *superior parietal cortices*), and the occipital lobe (left *lateral occipital*). For *g*_ii_, the bilateral *transverse temporal* cortex were involved through the training. The activations of the other lobes occurred only in session 3 before vanishing in session 4. They involved a reduced number of regions of the frontal (left *rostral middle frontal*), the parietal (left *inferior parietal* cortex and the right *posterior cingulate*), and occipital lobes (left *lateral occipital* cortex). The parameter *τ*_e_ again yielded the most extensive and consistent activation patterns across the lobes and training sessions. In the frontal lobe, the left *pars opercularis* was consistently involved until the last session. The other lobes showed consistent activations in all four sessions: in the parietal lobe (the *supramarginal* gyrus bilaterally), the *inferior parietal* cortex, the *superior parietal* gyrus, the *precuneus*, and the *posterior cingulate*); the temporal lobe (*transverse temporal*); and in the occipital lobes (the *cuneus* and the *pericalcarine* cortex). Moreover, the distribution of non-motor regions exhibiting significant activation shifted from 24 regions in session-1 to only 14 most relevant non-motor regions by session-4.

Overall, these findings revealed significant session-by-session differences in brain activity between MI and RS conditions. Notably, differential activation patterns were observed across key sensory, motor, and cognitive processing regions across sessions. Specifically, the parameters *g*_ei_ and *g*_ii_ exhibited lower values during RS compared to MI across sessions, whereas *τ*_e_ demonstrated a significant reduction in MI relative to RS for the activated regions. These results highlight the dynamic and task-dependent modulation of brain activation patterns in MI compared to RS.

Together, these MEG-derived session-wise changes, as illustrated in Fig. 5, indicate a learning-related refinement of cortical engagement, whereby early diffuse activation progressively converged onto task-relevant sensorimotor regions through region-specific modulation of neural population dynamics. Finally, the number of significantly activated regions (*p*_FDR_ *<* 0.05) declined from 29 in session-1 to 19 in session-4, reflecting a transition from diffuse to more targeted sensorimotor activation with training. This progressive reduction in the spatial extent of activation was robustly captured by the excitatory time constant *τ*_e_. Similar analyses were performed on EEG modeled data (refer to the supplementary document).

### Event-related desynchronization and synchronization are associated with excitatory time constant

Event-related desynchronization (ERD) and synchronization (ERS) during MI are characterized by distinct spectral signatures: ERD manifests as a reduction in alpha and beta frequency power, i.e., negative ERD/S (indicating regional activation), whereas ERS reflects sustained or increased oscillation, i.e., positive ERD/S (indicating inactivation) (26). To link these well-established physiological markers of MI to the model-derived parameters, we examined whether regions exhibiting ERD/S showed systematic changes in the excitatory time constant (*τ*_e_) and neural gains (*g*_ei_, *g*_ii_) during MI compared to RS. ERD/S was quantified from empirical spectra as the decibel (dB) change in band-limited *α* (8-12 Hz) and *β* (13-30 Hz) power between MI and RS 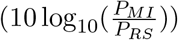, summarized using a composite *α* − *β* ERD/S measure.

First, to assess the global association between ERD/S and model-derived parameters in each session, we computed the relative change in *τ*_e_ for each region and subject as Δ*τ*_e_ (MI/RS-1) between MI and RS. Across regions, ERD/S exhibited a robust positive association with Δ*τ*_e_ for any given session (Spearman’s correlation *ρ* : 0.41 ≤ *ρ* ≤ 0.55; all four sessions, Bonferroni corrected *p <* 0.005) (Fig. 7-A), indicating a consistent global relationship between oscillatory power changes and excitatory population dynamics. Notably, regions that exhibited significant activation in our condition-effect analysis (Section 3.5) also exhibited desynchronization – ERD (negative ERD/S), whereas inactivated regions predominantly exhibited ERS (positive ERD/S), consistent with the well-established functional distinction between ERD and ERS. This is illustrated in Fig. 7-A, where activated regions are in the negative quadrant and inactivated regions are in the positive quadrant.

**Fig. 7.**
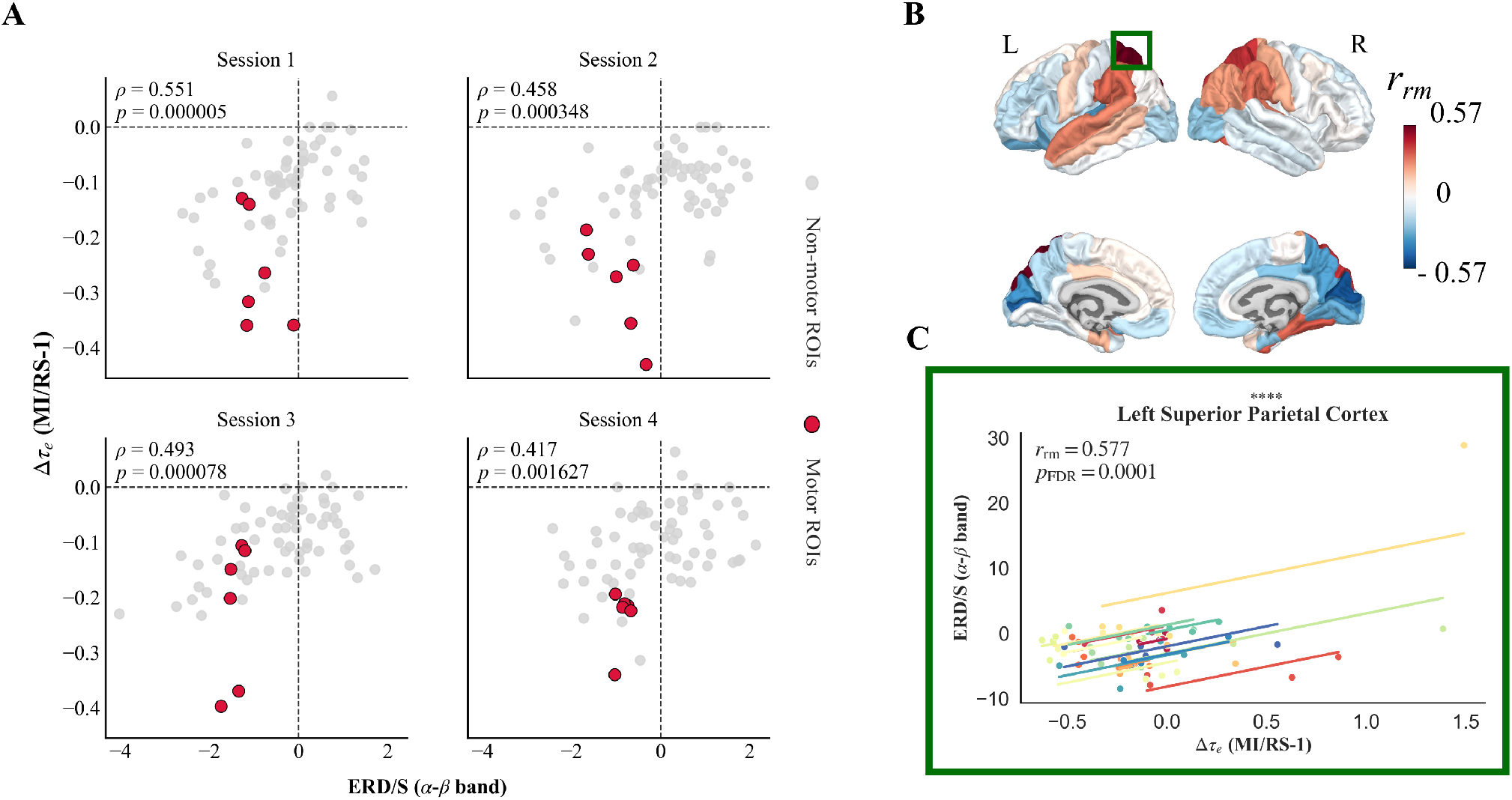
Significant association between excitatory time constant and ERD/S at the global and local level. A shows a significant positive association of ERD/S activity with the reduced excitatory time constant (Δ*τ*_e_) during MI at the global level in any given session (Spearman’s correlation *ρ* : 0.41 ≤ *ρ* ≤ 0.55, all sessions Bonferroni corrected *p <* 0.005). In each of the subplots, each point represents one ROI (median across subjects); motor regions are highlighted. Motor areas show consistent negative ERD/S (red) activity in all sessions, reflecting event-related desynchronization and task-related activation. Spatial map (B) shows variation in repeated measure correlation coefficient (*r*_*rm*_) between ERD/S and Δ*τ*_e_ within subjects, across sessions, computed for each of the 68 cortical regions. The map reveals both direction and magnitude of the association (*r*_*rm*_) for each region, with warm and cool colors indicating positive and negative association, respectively, which varies between −0.577 ≤ *r*_*rm*_ ≤ +0.577. The scatter plot (C) illustrates an important cortical region involved in MI – left superior parietal cortex, showing a significant positive within-subject association between ERD/S and Δ*τ*_e_ across sessions, where each line represents a subject-specific regression slope (*n* = 19,****: *p*_FDR_ *<* 0.0002, *r*_*rm*_ = 0.577). The robust positive correlation (*r*_*rm*_ = 0.577) indicates that stronger event-related desynchronization (reduction of *α* − *β* power) is consistently associated with larger reductions in *τ*_e_ during MI for that region.

Thereafter, we analyzed the spatial distribution of ERD/S for each of the 68 cortical regions to assess the coupling between longitudinal changes in oscillatory power (ERD/S) and excitatory population dynamics within a subject (intra-subject), after accounting for the non-independence of training sessions, reflecting training effect. Repeated-measure correlation analysis revealed a robust within-subject association between Δ*τ*_e_ and ERD/S in several cortical regions. The association varied regionally, from a negative to a positive association (−0.57 ≤ *r*_*rm*_ ≤ 0.57) as shown in the brain plot in Fig. 7-B. In particular, the left superior parietal cortex showed a significant positive association (*r*_*rm*_ = 0.577, *p*_FDR_ = 0.00014), which is known to be involved in motor-imagery and working memory. The scatter plot in Fig.7-C, shows session-to-session reductions in *τ*_e_ during MI consistently accompanied by stronger desynchronization (negative ERD/S) within individual subjects in the left superior parietal cortex. This significant positive association suggests that a reduction in excitatory time constant at the population level is associated with a reduction in *α* − *β* oscillatory power, and vice versa, in the left superior parietal cortex.

We next examined whether ERD/S was associated with changes in neural gains globally. ERD/S exhibited a significant negative association with the E/I coupling neural gain *g*_ei_ for any given session from session 1 to 3 (Spearman’s correlation *ρ* : −0.48 ≤ *ρ* ≤ −0.42, Bonferroni corrected *p <* 0.005). This negative association suggests that regions showing stronger desynchronization (more negative ERD/S change) are characterized by increased *g*_ei_, reflecting stronger recurrent interaction among E/I neuronal populations. Conversely, regions showing synchronization (positive ERD/S change) are characterized by a relative decrease in *g*_ei_. Importantly, this association weakened and lost significance by session 4 (Bonferroni corrected *p >* 0.05) (Fig. S5-A). The self-inhibitory neural feedback gain *g*_ii_ did not show any significant association with ERD/S (Fig. S5-B).

Together, these findings suggest that coordinated reconfigurations in E/I neuronal population dynamics underpin ERD/S during MI. Stronger desynchronization is potentially driven by faster excitatory postsynaptic population responses (reduced *τ*_e_). Thus, these associations position *τ*_e_ as a robust population-level correlate for MI task-related desynchronization.

## Discussion

This work represents the first biophysical modeling study in the field of BCI research to use a linear NMM for simulating motor task-related MI data, by modulating the scale-free spectral characteristics of the driving input to the excitatory/inhibitory neural populations. Using this novel modeling approach of mi-NMM to simulate MI versus RS, we successfully captured M/EEG spectra of individual brain regions during both conditions. Our findings demonstrate that this approach not only effectively models the MI and RS condition but also identifies key brain regions activated during MI across different BCI training sessions. In doing so, we provide new insights into the neural mechanisms underlying BCI training, paving the way for potential improved interpretability and optimization of BCI systems.

The mi-NMM we used here allows us to relate the model’s parameters to the neurophysiological functions of the cerebral cortex. This is especially true for the motor cortex, which was recorded by both MEG and EEG during the right-hand grasping motor imagery task. By fine-tuning these model parameter values within a biophysically realistic range, we were able to closely match the PSD shapes of the M/EEG and modeled spectra, while capturing the power difference between MI and RS. This allowed us to further investigate the link between the physiological mechanisms underpinning BCI training and the neurophysiologically meaningful model parameters.

While investigating the effect of MI compared to RS in each session, we analyzed regions of cerebral cortex based on DK atlas independently for each parameter, region, across individuals (inter-subject) in M/EEG. Here, we could observe that with an increase in session the mi-NMM parameters were able to specifically capture various important targeted regions with strong effect size, that are activated during the MI-BCI task, as shown in Fig. 5. Parameter E/I coupling neural gain (*g*_ei_), reflecting the functional interaction strength between the excitatory-inhibitory neuronal populations, showed significant changes in a limited, yet functionally critical areas in session 1. Specifically, significant regions that got activated as a result include the right inferior parietal cortex which is involved in sustaining attention over time (27); left lateral occipital cortex for visual memory recognition, visual object completion (28); left pars opercularis, and right superior parietal cortex essential for the manipulation of information in working memory and supports executive processes required to rearrange and mentally transform information (29). This initial pattern of activation might have served as a baseline or scaffold that enabled subsequent learning as we did not observe any significant regions in other sessions. As participants became more experienced with the MI-BCI task, the neural processes underlying sensory integration and motor planning likely became more efficient, possibly reducing the need for further modulation of the *g*_ei_ parameter. In other words, although *g*_ei_ was strongly engaged during the initial performance of the task, its influence appeared to stabilize with practice.

With the inhibitory neural gain parameter (*g*_ii_) we did not observe motor related or associated area getting activated at the beginning of the training. However, by session 3, increased functional engagement of task-relevant regions potentially reflect a shift in local population/circuit dynamics, where increased self-inhibitory neural feedback gain within the inhibitory neural population during MI reduced the overall inhibitory output, decreasing its net suppressive influence on excitatory neurons. This disinhibition potentially gave rise to net excitation with greater excitatory activity and allowed specific regions to become more functionally responsive during MI-BCI performance. Thus, involvement of these areas suggest that as learning progressed, inhibitory processes became more distributed across networks which included regions involved in sustained attention (inferior parietal and posterior cingulate) (27; 30); and in visual processing (lateral occipital) (28). Finally, by session 4, the pattern of activation associated with *g*_ii_ showed very distinct refinement, with significant activity observed in the right postcentral area. The involvement of this region might underscore the integration of somatosensory information and the fine-tuning of motor learning, while the continued significance of the left transverse temporal area from session 1 till the end reinforced its role in memory processing. This refined pattern in the final session potentially indicate that with sustained practice, inhibitory neural gain mechanisms evolved to support more specific functions—namely, enhancing voluntary motor control.

The excitatory time constant (*τ*_e_) at the population level represents the duration of excitatory post-synaptic responses. In our study, *τ*_e_ exhibited significant reduction during MI compared to RS (Fig. 5) across sessions for the activated regions. In session 1, *τ*_e_ related activity was widely distributed in many regions suggesting that at the beginning of BCI training, the duration of excitatory response are widely distributed, reflecting a generalized and random recruitment of neural circuits involved in sensory processing, higher-order cognition, and motor coordination. By session 2, as learning began, the activation became more focused to regions involved in visual processing and mental imagery (cuneus, and lingual gyrus) (31; 32; 33; 34), and error detection (left insula) (35). This indicates an early refinement of the timing of excitatory response, as the brain began to adapt to the temporal dynamics of the specific motor imagery condition (36). Then, additional areas seemed to get recruited such as the left inferior parietal and temporal regions, suggesting that, with continued practice, the duration of the excitatory neural population response fine-tuned to support more efficient sensorimotor integration and cognitive processing. Finally, in session 4, *τ*_e_ exhibited the most extensive and robust activation pattern, with significant involvement of regions crucial for motor imagery and motor hand tasks (precentral, paracentral and postcentral gyri) (37; 38); memory processing (precuneus, posterior cingulate, entorhinal cortex) (39; 30; 40); and sustained attention (inferior parietal and supramarginal regions) (27; 41).

Therefore, this progressive expansion and increased specialization of *τ*_e_ between sessions as shown in Fig. 5, potentially suggest that with repeated practice, the excitatory temporal profiles became finer tuned. The role of *τ*_e_ is also noteworthy because it showed a robust association with the MI-related ERD/S phenomenon in the alpha and beta frequency bands, as shown in Fig. 7-A. Physiologically, active processing places neurons in a high-conductance state, characterized by a reduction in effective membrane time constants due to increased synaptic drive and low resistance (42). Within the mi-NMM framework, this physiological regime potentially reflected at the neuronal population level as a faster decay of excitatory population responses (reduced *τ*_e_).

We also observed a decrease in the total number of activated regions from session 1 (29 regions) to session 4 (19 regions). This reduction potentially suggests that, with training, subjects were able to selectively engage the most relevant regions responsible for the MI task, thereby improving their control over the BCI. Such observations are in line with previous findings, where event-related desynchronization was observed as a significant power decrease in alpha and beta frequency bands (43). This was observed in cortical regions associated with key cognitive and sensorimotor functions relevant to BCI control, namely: sensorimotor integration and motor imagery (superior parietal lobule), decision-making and memory consolitation (middle anterior cingulate gyrus), behavioral adaptation (posterior ventral cingulate gyrus and frontomarginal gyrus), visual tracking (lingual gyrus), mental rotation and working memory (orbital part of the inferior frontal gyrus).

The EEG-fitted modeled parameters also evolved from session 1 through session 4 (Fig. S4). Initially, no specific brain regions were captured during the early training sessions. However, by the end of the training, in session 4, the parameters were significantly different in relevant regions for the MI-BCI task. This suggests that although subjects were initially naïve, training enabled them to engage the necessary brain regions for BCI. The E/I coupling neural gain parameter (*g*_ei_) showed no significant regional effects in session 1 but by session 4, it captured critical regions, including: Occipital lobe (bilateral cuneus, left lateral occipital)- involved in visual pattern detection, tracking visual motion, visual memory recognition, and visual mental imagery (39; 31; 32; 28); Parietal lobe (left inferior parietal)-associated with somatosensory processing and spatial awareness (44); Frontal-motor area (left precentral cortex) - responsible for motor control and execution (45). Similarly, with the inhibitory neural gain parameter (*g*_ii_), we observed some important regions (bilateral cuneus, left isthmus cingulate, left lingual, right parahippocampal) that are involved in spatial memory and navigation, memory formation, episodic memory retrieval, motor detection and tracking (46; 33; 34; 47; 48) in session 4. Lastly, the excitatory time constant (*τ*_e_) in session 4 showed significant condition effect in supramarginal gyrus, which is involved in somatosensory processing task (41; 49).

Thereafter, in order to further validate our observations, another MI-based EEG BCI dataset was brought for analysis. In this dataset, 15 subjects (RH, 8 F) performed a MI task to grasp an object using a robotic arm. They use a combination of gaze to indicate the position to reach and either performed MI of the right hand or resting state to grasp or not the object presented. They actively controlled the robot for 3 runs with 10 trials per condition. The protocols and results of the findings can be found in (50). Among the control strategies used, we only focused on the strategy where the robot moved prior to the MI task as it showed the highest level of desynchronization across subjects previously. In order to reduce variability of the findings due to the processing pipeline, we performed source reconstruction using wMNE and used parcellation with PCA over dipoles to get the Desikan-Killiany atlas in the different conditions of rest and MI. This set of subjects did not have their MRI scan meaning that the inverse method was done using fsaverage (an MRI template on the average of 40 subjects (51)). This means regions involved are to be interpreted cautiously as some of them could be overlapping over points specific to the different subjects.

The dataset revealed a decrease in PSD during MI with respect to RS in the broad range *α*-*β* bands, specifically in the post central region (52; 53) even though the mi-NMM parameters were not significant, contributing to confirm the MI of the right hand taking place in the different subjects. Furthermore, different regions that were mentioned in the previous analysis (54) have significantly different mi-NMM parameters in MI versus RS, shown in Fig. S6. Among them, we can pin out the caudal middle frontal region (55; 56) and the paracentral region (57) both used for motor planning to facilitate the execution of goal-directed actions and attention allocation; and the frontal pole associated with high-level cognitive functions like decision-making (with the limitation that it could be artifacted by gaze activity). The superior parietal lobe was also found to be activated. This region is found to be a central hub of multisensory integration and spatial perception (58), and is also involved in sense of agency and visuomotor coordination. These findings reinforce previous results on the impact of the robotic arm over subjects in their integration of the device to their mental workspace. Furthermore, our results based on the mi-NMM parameters in this validation dataset show that they can capture activation in the relevant regions as was the case with our main dataset. Other regions that were previously not listed (54) had significantly different mi-NMM parameters in MI – they may be relevant to performing MI such as the pericalcarine and cuneus regions involved in visual processing (59); and the medial orbitofrontal lobe associated with reward processing (60; 61) that may be relevant to the neurofeedback in MI.

Moreover, an important point of discussion in this regard is the reproducibility of the model findings, especially given that the experimental conditions and the tasks performed in the two BCI trainings are completely different. Despite these differences, mi-NMM consistently identified brain regions that overlap as statistically significant (*p*_FDR_ *<* 0.05) across the two datasets, suggesting robustness in its ability to capture potential neurobiological correlates. Specifically, in the case of the parameter *g*_ii_, two regions emerged as replicable in both datasets. First, the parahippocampal cortex (responsible for episodic memory formation and memory retrieval), and second, the Cuneus region (involved in visual detection of patterns, visual mental imagery and attention modulation). This overlap may reflect a shared visual or attentional demand across tasks, even if they differ in cognitive context.

Similarly to the previous dataset, condition effects observed with *g*_ii_ were associated with lower values in RS compared to MI, whereas *τ*_e_ exhibited lower values in MI than in RS. Notably, these consistencies between datasets—particularly in the absence of identical experimental conditions—demonstrate the potential generalizability of the mi-NMM and its capacity to identify relevant neurobiological correlates. However, not all parameters exhibited such reproducibility. For instance, *g*_ei_ and *τ*_e_ did not yield overlapping regions between datasets, suggesting that parameter-specific effects may be more sensitive to task or dataset characteristics.

Another key aspect to consider is whether using a linear instead of a non-linear model is justified in this case. The key advantage of using a linear model is that we can compute the PSD directly and then compare it with the real data. This reduces the computational load substantially and allows for a wider search of optimal model parameters. Biophysically, it is unclear which model captures the empirical features correctly. However, prior evidence suggests that a linear model may be sufficient in many cases. Most prominently, linear models outperformed non-linear models in predicting resting-state fMRI time series, owing to the linearizing effects of macroscopic neurodynamics and neuroimaging due to spatial and temporal averaging, observation noise, and high dimensionality (62). Lastly, network-level linear NMM of this kind outperformed a non-linear NMM in capturing MEG spectra accurately (18). While the model presented here is simpler than the nonlinear NMMs, one can ask if a simpler model than this linear NMM can be used for this study. From our knowledge, any model simpler than this linear NMM will not be able to capture the basic biophysical mechanism of the interaction between excitatory and inhibitory populations, which we aimed to investigate in this study. Therefore, a linear NMM of this kind is appropriate to test the hypothesis of this study.

In this regard, a critical aspect of our modeling approach is the driving input/stimulus, which we define as aggregate pre-synaptic input/background activity which is internal or arising from distant cortical/sub-cortical regions. In the RS condition, this background activity is considered to be largely asynchronous and weakly correlated across neuronal populations. Thus, under this condition it is considered to be white Gaussian noise, in line with prior studies. In contrast, during MI condition, we posit that these driving inputs are no longer purely asynchronous as reflected by the significant increase in the aperiodic spectral exponent values during MI (Fig. 2). Task engagement potentially recruits coordinated activity across functional networks mediated via structural connectivity across brain regions, leading to temporally structured and correlated afferent scale-free drive. Thus, to capture this regime, we explicitly modeled this as a parsimonious approximation of the temporally correlated drive, represented by colored noise with a power-law form of 1*/*(*f*^*k*^). Moreover, this modeling approach is neurophysiologically grounded in the framework of scale-free brain dynamics. Previous studies on scale-free brain activity (12; 13) demonstrated that the background arrhythmic activity of the brain, which is often treated as noise, exhibit a robust 1*/f* temporal structure that is functionally significant. According to these developments, scale-free brain activity in noninvasive recordings like MEG and EEG offer a window into the population level activity of cortical pyramidal neurons rather than being a typical noise, where the exponent of this scale-free activity is not static but is systematically modulated by task demands, reflecting shifts in brain’s functional operating point that facilitates information processing. Therefore, to capture this task-induced modulation in arrhythmic brain activity, we used colored noise for the MI task condition.

However, all the non-linear NMM (23; 24; 25; 16) as well as the limited modeling studies that have been done previously using non-linear NMMs to model motor tasks have attempted to capture task effects by adjusting the mean and variance of the white noise input (which is used to simulate RS). For instance, in (17), where the motor task involved grasping a pressure sensor with the right hand to measure grip strength, the mean and standard deviation of the Jansen-Rit model’s white noise parameter were adjusted to fit to the alpha band EEG PSD data. Similarly, in (16), where data were recorded during right-hand finger movements and the execution of a working memory task, the Wendling et al. (63) NMM was modified by incorporating white noise with a defined mean and variance. The mi-NMM consists of a linear NMM in which the noise parameter remains white noise with a constant mean and variance in RS. However, for the MI-BCI task, we modeled this input as colored noise. This is because unlike non-linear neural mass models, simply changing the mean and variance of the white noise will not change the shape of the PSD – it will only modify the amplitude. Therefore, we needed to modify the color of the noise in our case and our MI modeled spectra showed very less error rate compared to RS model, reflected in RMSE metric (Fig. 3-E, F) with both M/EEG modalities. Moreover, our empirical results (Section 3.1) Fig. 2 demonstrate significant task-dependent increase in aperiodic spectral exponent values during MI, supporting this hypothesis. By transitioning the input from white to colored noise, the mi-NMM attempts to capture a shift in scale-free brain activity from asynchronous background drive in RS to temporally structured task-driven input during MI.

Moreover, the current study consists of several limitations. In this study, the training sessions and number of participants are limited, and future studies replicated in larger and more diverse cohorts will help generalize the findings universally. Secondly, the fitted parameters were obtained using a stochastic global optimization procedure with one initial starting point, therefore a systemic multi-start optimization or posterior distribution characterization will help establish reliability of these estimated parameters. Additionally, every region is fitted to the model separately and using a whole-brain model that incorporates the structural connectivity can potentially be used to explain network-level reorganization during learning.

Interestingly, observation from our analysis of EEG and MEG modeled data reveals that in EEG, *τ*_e_ does not show any condition effect or capture any region except one in session 4, making it difficult to pinpoint specific areas of activity for this parameter. However, in MEG, *τ*_e_ reliably highlights regions within the cerebral cortex during all four sessions, suggesting that MEG provides a more stable and regionally specific representation of neural dynamics compared to EEG in this case. Similar to the findings in our previous study (43), a greater number of regions of interest exhibited significant condition effects in MEG compared to EEG. This discrepancy can be attributed to the distinct biophysical properties of the two modalities. EEG signals are attenuated by the varying conductivities of intervening tissues, such as the scalp, skull, and cerebrospinal fluid (64), and are sensitive to both tangential and radial components of cortical dipoles. In contrast, MEG offers superior spatial resolution and is primarily sensitive to the tangential components of neural sources (65). Consequently, EEG and MEG do not necessarily capture the same aspects of brain activity. Furthermore, participants were instructed to lean their heads back within the MEG helmet during data acquisition. This positioning brought the MEG sensors closer to the parietal and occipital regions, potentially enhancing the sensitivity to posterior brain activity relative to signals arising from the frontal areas. Lastly, this is a correlation-based study and cannot be directly used to infer causual mechanisms that require controlled experiments.

In conclusion, this study shows that mi-NMM derived from biophysically interpretable parameters capturing underlying E/I dynamics are capable of replicating task dynamics across sessions in a way that closely mirrors real data. Its ability to track session-to-session variations provides compelling evidence that E/I coupling may play a role as an underlying brain mechanism during BCI learning. Looking ahead, these insights open the door to integrating E/I coupling into the BCI framework, as it emerges as a relevant feature in shaping and informing the training process. More broadly, embedding these mechanisms into BCI design could contribute to more refined and neurobiologically grounded models, ultimately enriching both the theoretical and practical aspects of BCI development.

## Methods

### Brain-Computer Interface data collection and analysis

In this study, we worked with a dataset composed of a cohort of 19 healthy right-handed subjects (mean age:27.4 ± 4.0 years; 12 men) participated in a BCI study. All participants were BCI-naive, healthy and free from medical or psychological disorders. This study was conducted in strict accordance with the principles outlined in the Declaration of Helsinki and it was not a clinical trial. Following a thorough explanation of the study protocol, all participants gave written informed consent to take part in the experimental procedures, and the study was approved by the ethical committee CPP-IDF-VI of Paris (no. 2016-A00626-45).

Subjects underwent a two-week motor imagery-based BCI training program consisting of four sessions. At the beginning of each session, the subjects underwent a 3-minute RS with eyes opened. The BCI experiments consisted of a standard one-dimensional, two-target box task (66), where subjects controlled the vertical position of a cursor moving at a constant velocity from left to right to reach a target displayed on the right part of the screen. To reach the up target, the subjects were instructed to perform sustained right-hand motor imagery (MI) by modulating neural activity in the alpha (8–12 Hz) and beta (13–30 Hz) frequency bands. To hit the down target, the subjects had to remain at rest–which we refer here as baseline condition. Each of the six runs consisted of 32 trials, distributed equally and randomly among the targets.

Neural activity was recorded using simultaneous EEG and MEG modalities with a 74-channel EEG system (Ag/AgCl passive sensors, Easycap, Germany) following the 10-10 montage. MEG data were acquired using 102 magnetometers and 204 gradiometers (Elekta Neuromag TRIUX) in a magnetically shielded environment. The signals were sampled at 1 kHz (0.01–300 Hz bandwidth) and then downsampled to 250 Hz. MEG signals were preprocessed with temporal Signal Space Separation using MaxFilter (67). Eye and cardiac artifacts were removed from EEG/MEG signals via Independent Component Analysis with the Infomax approach in FieldTrip (68; 69). EEG electrodes T9 and T10 were excluded from further analysis. Source reconstruction was performed using the Boundary Element Method (70; 71) with three-layer MRI-derived surfaces (scalp, inner skull, outer skull; 1922 vertices each). The weighted Minimum Norm Estimate (72; 73; 74) was used to estimate cortical sources via Brainstorm (75), with an identity noise covariance matrix. The dipole orientations were constrained to the cortex. Individual source estimates were projected onto the MNI-ICBM152 template (76) using Shepard’s interpolation.

Power spectra were calculated for each subject, session, and condition in the source space using the Welch method (1 s window, 50% overlap). For the rest condition, the estimation was performed directly from the 3-minute recording. For the MI condition, the power spectra were computed from the feedback period (t = 3–6 s) and averaged across trials. To localize the anatomical structures associated with the obtained clusters, we used the Desikan killiany (DK) atlas (77) without restricting the analysis to the motor or sensorimotor regions. The dataset was structured as (19 × 68 × 126) per session and condition, where 19 represents subjects, 68 corresponds to DK atlas-defined brain regions, and 126 represents frequency bins used in spectral analysis.

For more information on the experiment and the data processing, please refer to (43) and raw data is available here (78).

### Model description

We used a linearized NMM (18; 19; 20) to extract excitatory and inhibitory neural subpopulation parameters. The model at the mesoscopic level (regional model), for every region *k* (*k* varies from 1 to *N* and *N* is the total number of regions) based on the DK parcellation, is modeled as the sum of excitatory signals *x*_e_(*t*) and inhibitory signals *x*_i_(*t*). Both excitatory and inhibitory signal dynamics consist of a decay of the individual signals with a fixed neural gain, incoming signals from populations that couple the excitatory and inhibitory signals, and driving input as Gaussian white noise for resting state and colored noise for mentally active state. The model equations for the excitatory and inhibitory signals for every region of the DK atlas that describes the rate of change of signal or firing rate in excitatory and inhibitory neuronal populations are:

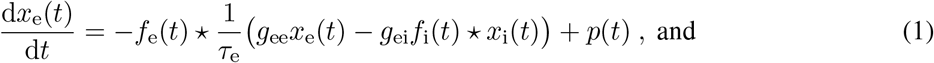

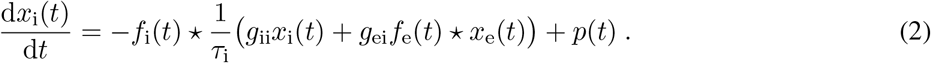

The symbols used in the equations are as follows: ⋆ denotes convolution; parameters *g*_ee_, *g*_ii_, and *g*_ei_ represent neural gains for excitatory, inhibitory and for coupling neural gain between excitatory and inhibitory populations, respectively; *τ*_e_ and *τ*_i_ are the time constants of excitatory and inhibitory populations, respectively; *p*(*t*) is driving input which is presynaptic input/stimulus; *f*_e_(*t*) and *f*_i_(*t*) are average neural impulse response functions. Parameters *g*_ei_, *g*_ii_, *τ*_e_, and *τ*_i_ were estimated for each region of interest (ROI), while *g*_ee_ was fixed at 1 for parameter identifiability. The excitatory and inhibitory time constants characterize the duration of post-synaptic responses (modeled by a gamma-shaped function) in each neuronal populations. They also determine the rate at which a local signal dissipates in the absence of other inputs. A lower time constant indicates a faster decay of these responses, whereas a higher time constant corresponds to a slower, long-lasting decay. The neural gain parameters: excitatory (*g*_ee_) and inhibitory (*g*_ii_) neural gain represent the strength of self-excitatory and self-inhibitory feedback gain within their own respective neuronal populations, and the coupling excitatory-inhibitory (E/I) neural gain (*g*_ei_) reflects the strength of interaction between the two populations, as in how one population influences the activity of the other.

We can get a closed-form solution in the Fourier domain since the two equations above are linear, as shown below. The Fourier transform *F*() is taken at an angular frequency *ω*, which is equal to 2*πf*, where *f* is the frequency in Hz. The Fourier transform of equations 1 and 2 will be the following, where *F*(*x*_e_(*t*)) = *X*_e_(*ω*) and *F*(*x*_i_(*t*)) = *X*_i_(*ω*), and j is the imaginary unit.

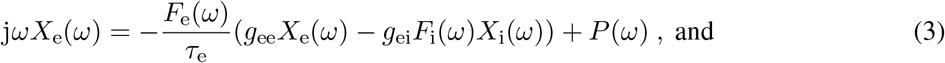

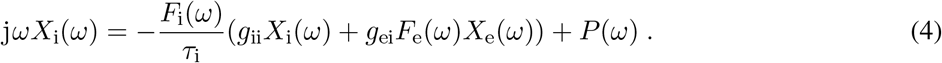

On solving the above equations 3 and 4, *X*_e_(*ω*) and *X*_i_(*ω*) are:

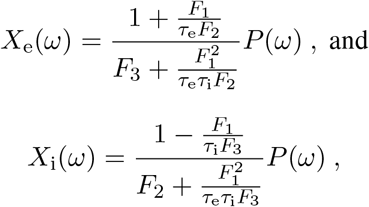

Where

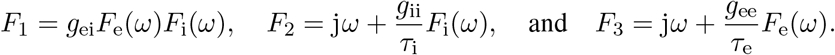

Then, the transfer functions *H*_e_(*ω*) and *H*_i_(*ω*) can be separated, and we get:

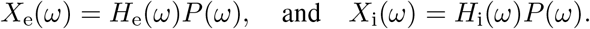

Thus, for the total neural population at each region:

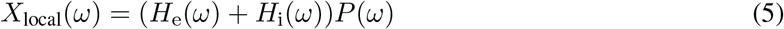

The driving input *p*(*t*) is characterized by its power spectral density (PSD) *S*_*p*_(*ω*), which is assumed to be identical for excitatory and inhibitory local populations within each region. To simulate RS condition, *p*(*t*) was modeled as white noise, characterized by frequency-independent constant PSD. To simulate the MI condition, *p*(*t*) was modeled as frequency-dependent, colored noise with a power-law spectrum, such that *S*_*p*_(*ω*) ∝ 1*/f* ^*k*^, where *k* ∈ [0, 1].

Given the linear time-invariant nature of the model, the simulated output PSD was obtained for each region as:

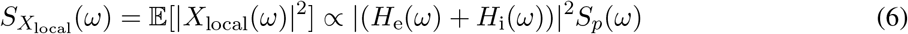

where E[·] denotes expectation. The resulting spectra were expressed in dB scale as 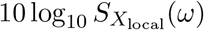. Thus, overall in the end we are only interested in the PSD of the signal. Although the equations include a stochastic input term, we do not solve them in the time domain as stochastic differential equations. Our analysis is restricted to second-order statistics of the system output, which can be obtained directly in the frequency domain under assumptions of linearity and stationarity. Hence, the spectral properties of *p*(*t*) are defined in-terms of PSD, obtained from second-order frequency domain statistics as *S*_*p*_(*ω*) ∝ E[|*P*(*ω*)|^2^], where *P*(*ω*) is Fourier transform of finite-length realization of the input process. Accordingly, our analysis focuses exclusively on PSDs, which are deterministic quantities characterizing the second-order structure of the stochastic signals.

### Model parameter estimation

Model parameters were estimated for each region, each subject, and across all sessions, for frequency range of [3-30] Hz. For every subject, each region’s spectrum was modeled using equations 3 and 4, and finally the total spectrum PSD obtained from equation 6 was compared to the empirical spectrum. The goodness of fit of the model was evaluated by calculating Pearson’s correlation coefficient between the modeled power spectra and the empirical source-localized (M/EEG) spectra. This goodness-of-fit value was used to optimize the model parameters.

All the model parameters were set within a biophysically realistic range based on prior experiments (18), and the parameter optimization was performed using the basin-hopping global optimization algorithm in Python (79). The initial guess for all parameters and the parameter bounds used for estimating the modeled spectra are provided in Table 1. To ensure model stability, we defined five different bounds for the excitatory-inhibitory coupling neural gain (*g*_ei_) (80). Initially, a larger bound was applied to *g*_ei_, and if the optimal model parameter crossed the stability boundary, the optimization was repeated with a smaller bound. This process was iterated up to five times to ensure that the final optimal parameters corresponded to stable model solutions. The model parameters and their bounds are specified in Table 1. In addition to these parameters, the exponent of the colored input noise was also optimized for in the case of MI, within the range of [0,1] as mentioned in the previous section. Importantly, all the intrinsic model parameters (*g*_ei_, *g*_ii_, *τ*_e_, *τ*_i_) were estimated independently for each region, subject, session, and condition, and were allowed to vary freely within their respective bounds during optimization to maximize the goodness of fit between modeled and empirical spectra.

**Table 1.**
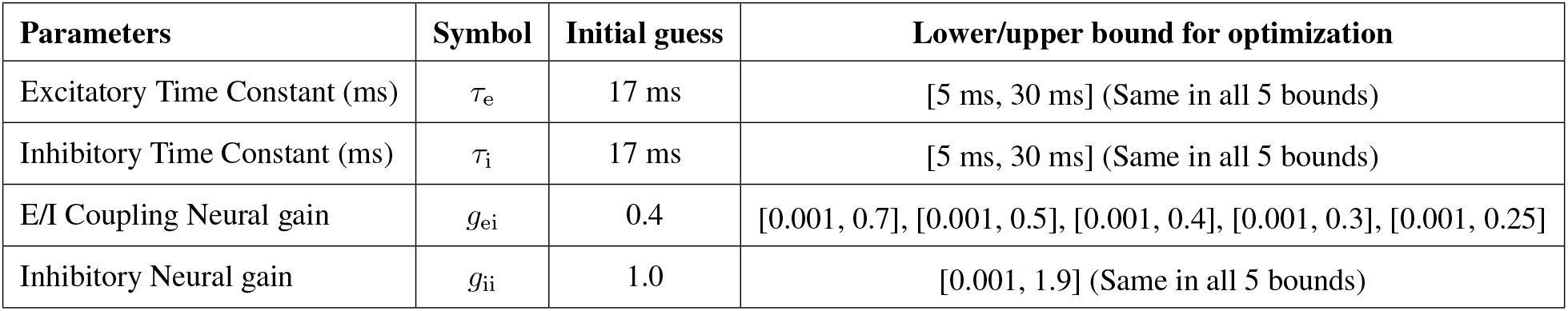
Model parameters with biophysically realistic initial guesses, and bounds for estimating the spectra.

The algorithm’s hyperparameters included the number of iterations (*niter*), temperature (*T*), and step size (*stepsize*), which were set to 5000, 0.1, and 4, respectively. If any parameter hit the specified bounds, the optimization was repeated with a step size of 6 for that specific ROI. Finally, the set of parameters yielding the highest Pearson’s correlation coefficient was selected. The cost function for optimization was defined as the negative Pearson’s correlation coefficient between the source-localized (E/MEG) spectra in the dB scale and the model power spectral density in the dB scale.

As this is the first pilot study, we started with comparing the most contrasting conditions. Therefore, we focused on the resting state recorded before the BCI experiment was started, and the MI performed during the BCI training. Specifically, we did not include the baseline condition in which subjects were instructed to remain at rest during the BCI training – the baseline condition may not correspond to strictly being in the resting state as the subjects can still continue to think about the experiment/their performance.

### Evaluation of modeled spectra & Parameters contribution

To evaluate the model performance, all the modeled spectra after optimization were evaluated using the root-mean square error (RMSE) computed between the modeled PSD and the empirical M/EEG PSD (both Z-score normalized), in addition to the Pearson correlation coefficient. Thereafter, to quantify the influence of individual model parameters and input exponent on the generated modeled spectra, we performed a local sensitivity analysis across all ROIs, subjects, and sessions for both RS and MI condition, in M/EEG imaging modalities. The sensitivity of each model parameter/exponent was quantified using a normalized perturbation-based finite-difference approach. Each parameter/exponent (*θ*) was independently perturbed one-at-a-time while keeping others fixed. For each parameter *θ*, the perturbation was defined as Δ*θ* = *ϵθ*, where an example *ϵ* = 0.10 corresponds to a relative perturbation of ±10%. For the modeled PSD, *S*(*ω*), the relative spectral sensitivity coefficient of the parameter *θ* was computed as,

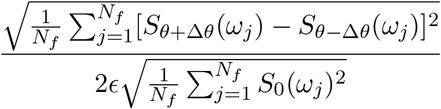

where *S*_*θ*+Δ*θ*_(*ω*_*j*_) and *S*_*θ−*Δ*θ*_(*ω*_*j*_) denote the modeled PSD at the *j*-th frequency bin (*N*_*f*_ is number of frequency bins), after increasing and decreasing parameter *θ* by a relative perturbation of *ϵ*, respectively and perturbation was done for multiple epsilon values (*ϵ* = 0.05, 0.10, 0.20, 0.30, 0.40). Here, *S*_0_(*ω*_*j*_) denotes the baseline modeled PSD generated by the optimized parameter values. Each parameter was perturbed independently, and we report the mean and standard error of the relative spectral sensitivity coefficients across all ROIs, subjects, and sessions for each parameter.

Similarly, to assess the influence of model parameters and input exponent on the modeled spectral slope, we performed a perturbation-based sensitivity analysis on each parameter/exponent across all ROIs, subjects, and sessions for MI condition. For each ROI, subject, and session, the optimized parameters and exponent was used to generate the baseline modeled spectrum, from which the baseline spectral slope (*m*_0_) was estimated. Each parameter/exponent (*θ*) was then independently perturbed one-at-a-time by *ϵ* while keeping all other parameters fixed and repeated for multiple *ϵ* (= 0.10, 0.20, 0.30, 0.40, 0.50) values. Then the influence of each parameter on the model spectral slope was quantified using a relative spectral slope sensitivity coefficient as,

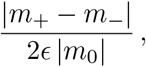

where *m*_+_ and *m*_*−*_ denotes the slopes of the perturbed spectra *S*_*θ*+Δ*θ*_(*ω*) and *S*_*θ−*Δ*θ*_(*ω*), respectively, that were generated following positive and negative perturbations, respectively. A larger slope sensitivity coefficient indicates greater sensitivity of the modeled spectral slope to the corresponding parameter, whereas a smaller value indicates that the spectral slope is relatively robust to perturbations of that parameter. All the slopes were calculated using the FOOOF (22) python package.

### Statistical analyses

To quantify the difference between the spectral exponent values between MI and RS, we used Paired t-Test and set significant threshold at 0.05, and to quantify the effects size of Paired-t Test we calculated the Cohen’s *d*_*z*_ as the mean of paired difference (MI – RS) divided by the standard deviation of the paired difference. Then to examine the potential condition related modulation between MI and RS in each session separately, we performed Wilcoxon signed rank test (81) on all four intrinsic model parameters across 19 subjects. We chose this non-parametric Wilcoxon test as the model derived biophysical parameters were not normally distributed. This analysis was conducted for each ROI to compare MI and RS conditions for all four parameters. For each session, we obtained 68 p-values (one for each brain region based on the DK atlas) per parameter, thus four sets of 68 p-values per session (one set for each parameter). Thereafter, we applied the false discovery rate (FDR) correction (82) to each of the four sets to account for multiple comparisons. We set a significance threshold at *p*_FDR_ *<* 0.05 to identify statistically significant regions for these parameters individually. The brain regions that met this significance criterion were identified for each parameter separately in each session. We used the same procedure both for EEG and MEG. Additionally, Wilcoxon effect size was computed by dividing the standardized Wilcoxon statistics (Z) with the square root of total number of sample size (N) in each session. Alongside, to specifically analyze the effect of MI in all left motor cortex areas in each session, we applied Wilcoxon signed rank-test and corrected for multiple comparisons using Bonferroni correction methods and set significant threshold at 0.05 and calculated the Wilcoxon effect size like the previous one.

Next, we analyzed how the session affected MI and RS in both M/EEG. We used the Friedman test (83) on the statistically significant left motor regions for the corresponding parameter from the session 4 condition effect analysis. Then, we corrected the two p-values (as two regions in left motor area in MEG) using the Bonferroni correction method and the effect size of the Friedman test, Kendall’s coefficient of concordance (W) was computed using Pingouin Python package.

Thereafter, we examined session-wise relationship between excitatory time constant (*τ*_e_) and group-level BCI performance using linear mixed-effects models (LMM) (84). Here, the BCI score is set as dependent variable, regional excitatory time constant (*τ*_e_) as a fixed effect, session as a fixed categorical factor, and subject as a random intercept to account for repeated measurements within subjects across sessions (Model: Score ~ *τ*_e_ + Session + (1 | Subject)). LMM was fitted independently for each of the 68 regions and FDR correction was applied for multiple comparisons, and implemented using statsmodels Python package.

Furthermore, to relate the model parameters to event-related desynchronization (ERD) and synchronization (ERS) dynamics, we conducted Spearman’s rank correlation to test the correlation between ERD/S and relative parameter (MI/RS-1) at the global level across 68 regions for each session and four session-specific *p* values were corrected using the Bonferroni correction by setting the significance at 0.05. Repeated measure correlation was performed to assess within subject coupling between Δ*τ*_e_ and ERD/S across sessions, accounting for the non-independence of repeated measurements across sessions. Analyses were performed for each of the 68 regions, and the resulting *p* values were corrected for multiple comparisons using FDR correction by setting the significance at 0.05.

## Data Availability

Minimally processed data is available in Networks for BCI (NETBCI) dataset: https://doi.org/10.57745/RBJRC7

## Code Availability

All the custom codes related to the paper can be found on this Github page: https://github.com/ApurbaApd/mi-NMM_bci_modeling

## Acknowledgements

Not applicable.

## Funding

The authors acknowledge support from Transatlantic Research Partnership from the FACE Foundation and Alzheimer’s Association grant AARFD-22-923931 and support from the European Research Council (ERC), Grant Agreement No. 864729.

## Author Contributions

P.V. and M-C.C. designed the study. A.D. analyzed the data, generated all the results, did subsequent analysis, and wrote the original draft of the paper. M-C.C. and T.V. preprocessed the data. All authors contributed to manuscript preparation. A.D., M-C.C., and P.V. reviewed, edited and revised the paper.

## Competing Interests

The authors declare no competing financial or non-financial interests.

## Additional information

Please see the supplementary document for the additional information.

## Notes

### Competing Interest Statement

The authors have declared no competing interest.

### Summary of Updates

Paper has been fully revised with a change in orders.

## References

[1] M. Clerc, L. Bougrain, and F. Lotte, Brain-Computer Interfaces 1: Methods and Perspectives. Wiley, wiley ed., 2016.

[2] M. Clerc, L. Bougrain, and F. Lotte, Brain-Computer Interfaces 2: Technology and Applications. Wiley, wiley ed., 2016.

[3] J. J. Shih, D. J. Krusienski, and J. R. Wolpaw, “Brain-Computer Interfaces in Medicine,” Mayo Clinic Proceedings, vol. 87, pp. 268–279, Mar. 2012.

[4] B. Z. Allison and C. Neuper, “Could anyone use a BCI?,” in Brain-Computer Interfaces (D. S. Tan and A. Nijholt, eds.), Human-Computer Interaction Series, pp. 35–54, Springer London, 2010.

[5] F. Lotte, L. Bougrain, A. Cichocki, M. Clerc, M. Congedo, A. Rakotomamonjy, and F. Yger, “A review of classification algorithms for eeg-based brain-computer interfaces: a 10 year update,” Journal of Neural Engineering, vol. 15, no. 3, p. 031005, 2018.

[6] M. Ahn and S. C. Jun, “Performance variation in motor imagery brain–computer interface: A brief review,” Journal of Neuroscience Methods, vol. 243, pp. 103–110, Mar. 2015.

[7] O. Alkoby, A. Abu-Rmileh, O. Shriki, and D. Todder, “Can We Predict Who Will Respond to Neurofeedback? A Review of the Inefficacy Problem and Existing Predictors for Successful EEG Neurofeedback Learning,” Neuroscience, vol. 378, pp. 155–164, May 2018.

[8] S. C. Kleih, F. Nijboer, S. Halder, and A. Kübler, “Motivation modulates the p300 amplitude during brain-computer interface use,” Clinical Neurophysiology, vol. 121, no. 7, pp. 1023–1031, 2010.

[9] C. Jeunet, E. Jahanpour, and F. Lotte, “Why standard brain-computer interface (bci) training protocols should be changed: an experimental study,” Journal of Neural Engineering, vol. 13, no. 3, p. 036024, 2016.

[10] R. Sitaram, T. Ros, L. Stoeckel, S. Haller, F. Scharnowski, J. Lewis-Peacock, N. Weiskopf, M. L. Blefari, M. Rana, E. Oblak, N. Birbaumer, and J. Sulzer, “Closed-loop brain training: the science of neurofeedback,” Nature Reviews Neuroscience, vol. 18, no. 2, pp. 86–100, 2017.

[11] D. S. Bassett, N. F. Wymbs, M. A. Porter, P. J. Mucha, J. M. Carlson, and S. T. Grafton, “Dynamic reconfiguration of human brain networks during learning,” Proceedings of the National Academy of Sciences of the United States of America, vol. 108, no. 18, pp. 7641–7646, 2011.

[12] B. J. He, J. M. Zempel, A. Z. Snyder, and M. E. Raichle, “The temporal structures and functional significance of scale-free brain activity,” Neuron, vol. 66, no. 3, pp. 353–369, 2010.

[13] B. J. He, “Scale-free brain activity: past, present, and future,” Trends in Cognitive Sciences, vol. 18, no. 9, pp. 480–487, 2014.

[14] T. Donoghue, “A systematic review of aperiodic neural activity in clinical investigations,” European Journal of Neuroscience, vol. 62, no. 7, p. e70255, 2025.

[15] M. Breakspear, “Dynamic models of large-scale brain activity,” Nature neuroscience, vol. 20, no. 3, pp. 340–352, 2017.

[16] M. Zavaglia, L. Astolfi, F. Babiloni, and M. Ursino, “A neural mass model for the simulation of cortical activity estimated from high resolution eeg during cognitive or motor tasks,” Journal of Neuroscience Methods, vol. 157, no. 2, pp. 317–329, 2006.

[17] Y. Zhang, Z. Li, H. Xu, Z. Song, P. Xie, P. Wei, and G. Zhao, “Neural mass modeling in the cortical motor area and the mechanism of alpha rhythm changes,” Sensors (Basel, Switzerland), vol. 25, no. 1, p. 56, 2024.

[18] A. Raj, C. Cai, X. Xie, E. Palacios, J. Owen, P. Mukherjee, and S. Nagarajan, “Spectral graph theory of brain oscillations,” Human Brain Mapping, vol. 41, no. 11, pp. 2980–2998, 2020.

[19] P. Verma, S. Nagarajan, and A. Raj, “Spectral graph theory of brain oscillations–revisited and improved,” NeuroImage, vol. 249, p. 118919, 2022.

[20] K. G. Ranasinghe, P. Verma, C. Cai, X. Xie, K. Kudo, X. Gao, H. Lerner, D. Mizuiri, A. Strom, L. Iaccarino, R. La Joie, B. L. Miller, M. L. Gorno-Tempini, K. P. Rankin, W. J. Jagust, K. Vossel, G. D. Rabinovici, A. Raj, and S. S. Nagarajan, “Altered excitatory and inhibitory neuronal subpopulation parameters are distinctly associated with tau and amyloid in alzheimer’s disease,” eLife, vol. 11, p. e77850, 2022.

[21] J. M. Fan, K. Kudo, P. Verma, K. G. Ranasinghe, H. Morise, A. M. Findlay, K. Vossel, H. E. Kirsch, A. Raj, A. D. Krystal, et al., “Cortical synchrony and information flow during transition from wakefulness to light non-rapid eye movement sleep,” Journal of Neuroscience, vol. 43, no. 48, pp. 8157–8171, 2023.

[22] T. Donoghue, M. Haller, E. J. Peterson, P. Varma, P. Sebastian, R. Gao, T. Noto, A. H. Lara, J. D. Wallis, R. T. Knight, et al., “Parameterizing neural power spectra into periodic and aperiodic components,” Nature neuroscience, vol. 23, no. 12, pp. 1655–1665, 2020.

[23] B. H. Jansen, G. Zouridakis, and M. E. Brandt, “A neurophysiologically-based mathematical model of flash visual evoked potentials,” Biological cybernetics, vol. 68, pp. 275–283, 1993.

[24] B. H. Jansen and V. G. Rit, “Electroencephalogram and visual evoked potential generation in a mathematical model of coupled cortical columns,” Biological cybernetics, vol. 73, no. 4, pp. 357–366, 1995.

[25] O. David and K. J. Friston, “A neural mass model for meg/eeg: coupling and neuronal dynamics,” NeuroImage, vol. 20, no. 3, pp. 1743–1755, 2003.

[26] G. Pfurtscheller and F. H. Lopes da Silva, “Event-related eeg/meg synchronization and desynchronization: basic principles,” Clinical Neurophysiology, vol. 110, no. 11, pp. 1842–1857, 1999.

[27] M. Husain and P. Nachev, “Space and the parietal cortex,” Trends in Cognitive Sciences, vol. 11, no. 1, pp. 30–36, 2007.

[28] Y. Lerner, T. Hendler, D. Ben-Bashat, M. Harel, and R. Malach, “Object-completion effects in the human lateral occipital complex,” Cerebral Cortex, vol. 12, no. 2, pp. 163–177, 2002.

[29] M. Koenigs, A. K. Barbey, B. R. Postle, and J. Grafman, “Superior parietal cortex is critical for the manipulation of information in working memory,” The Journal of Neuroscience, vol. 29, no. 47, pp. 14980–14986, 2009.

[30] R. Leech and D. J. Sharp, “The role of the posterior cingulate cortex in cognition and disease,” Brain, vol. 137, no. Pt 1, pp. 12–32, 2014.

[31] G. Plomp, A. Hervais-Adelman, L. Astolfi, and C. M. Michel, “Functional specialization and dynamic resource allocation in visual cortex,” Human Brain Mapping, vol. 31, no. 1, pp. 1–13, 2010.

[32] A. Qureshy, R. Kawashima, M. B. Imran, M. Sugiura, R. Goto, K. Okada, K. Inoue, K. Horie, S. Yoshioka, and H. Fukuda, “Functional mapping of human brain in olfactory processing: a pet study,” Journal of Neurophysiology, vol. 84, no. 3, pp. 1656–1666, 2000.

[33] Y. Kamitani and F. Tong, “Decoding seen and attended motion directions from activity in the human visual cortex,” Current Biology, vol. 16, no. 11, pp. 1096–1102, 2006.

[34] T. D. Waberski, R. Gobbelé, K. Lamberty, H. Buchner, J. C. Marshall, and G. R. Fink, “Timing of visuo-spatial information processing: electrical source imaging related to line bisection judgements,” Neuropsychologia, vol. 46, no. 5, pp. 1201–1210, 2008.

[35] D. Borsook, R. Veggeberg, N. Erpelding, R. Borra, C. Linnman, R. Burstein, and L. Becerra, “The insula: A “hub of activity” in migraine,” The Neuroscientist, vol. 22, no. 6, pp. 632–652, 2016.

[36] Y. Takenaka, T. Suzuki, and K. Sugawara, “Time course effect of corticospinal excitability for motor imagery,” European Journal of Neuroscience, vol. 54, pp. 6123–6134, 2021.

[37] T. A. Yousry, U. D. Schmid, H. Alkadhi, D. Schmidt, A. Peraud, A. Buettner, and P. Winkler, “Localization of the motor hand area to a knob on the precentral gyrus. a new landmark,” Brain: A Journal of Neurology, vol. 120, no. Pt 1, pp. 141–157, 1997.

[38] C. A. Porro, V. Cettolo, M. P. Francescato, and P. Baraldi, “Ipsilateral involvement of primary motor cortex during motor imagery,” The European Journal of Neuroscience, vol. 12, no. 8, pp. 3059–3063, 2000.

[39] B. N. Lundstrom, M. Ingvar, and K. M. Petersson, “Isolating the retrieval of imagined pictures during episodic memory: activation of the left precuneus and left prefrontal cortex,” NeuroImage, vol. 20, no. 4, pp. 1934–1943, 2003.

[40] A. D. Garcia and E. A. Buffalo, “Anatomy and function of the primate entorhinal cortex,” Annual Review of Vision Science, vol. 6, pp. 411–432, 2020.

[41] R. J. Perry and S. Zeki, “The neurology of saccades and covert shifts in spatial attention: an event-related fmri study,” Brain: A Journal of Neurology, vol. 123, no. Pt 11, pp. 2273–2288, 2000.

[42] A. Destexhe, M. Rudolph, and D. Paré, “High-conductance state of neocortical neurons in vivo,” Nature Reviews Neuroscience, vol. 4, no. 9, pp. 739–751, 2003.

[43] M. C. Corsi, M. Chavez, D. Schwartz, N. George, L. Hugueville, A. E. Kahn, S. Dupont, D. S. Bassett, and F. De Vico Fallani, “Functional disconnection of associative cortical areas predicts performance during bci training,” NeuroImage, vol. 209, p. 116500, 2020.

[44] J. Wang, Y. Yang, L. Fan, C. Xu, P. T. Fox, S. B. Eickhoff, C. Yu, and T. Jiang, “Correspondent functional topography of the human left inferior parietal lobule at rest and under task revealed using resting-state fmri and coactivation based parcellation,” Human Brain Mapping, vol. 38, no. 3, pp. 1659–1675, 2017.

[45] S. Rose, A. Guzzetta, K. Pannek, R. Boyd, M. Bastiani, L. Emsell, C. Falconer, S. Fiori, and D. K. Jones, “Structural hemispheric asymmetries in the human precentral gyrus hand representation,” Neuroscience, vol. 210, pp. 211–221, 2012.

[46] E. T. Rolls, “The cingulate cortex and limbic systems for emotion, action, and memory,” Brain Structure and Function, vol. 224, no. 9, pp. 3001–3018, 2019.

[47] A. M. Owen, B. Milner, M. Petrides, and A. C. Evans, “A specific role for the right parahippocampal gyrus in the retrieval of object-location: a positron emission tomography study,” Journal of Cognitive Neuroscience, vol. 8, no. 6, pp. 588–602, 1996.

[48] V. D. Bohbot, M. Kalina, K. Stepankova, N. Spackova, M. Petrides, and L. Nadel, “Memory deficits characterized by patterns of lesions to the hippocampus and parahippocampal cortex,” Annals of the New York Academy of Sciences, vol. 911, pp. 355–368, 2000.

[49] E. Ben-Shabat, F. Matyas, G. S. Pell, A. Brodtmann, and L. M. Carey, “The right supramarginal gyrus is important for proprioception in healthy and stroke-affected participants: A functional mri study,” Frontiers in Neurology, vol. 6, p. 248, 2015.

[50] T. Venot, A. Desbois, M. C. Corsi, L. Hugueville, L. Saint-Bauzel, and F. De Vico Fallani, “Intentional binding for noninvasive BCI control,” Journal of Neural Engineering, vol. 21, no. 4, 2024-07-25.

[51] B. Fischl, M. I. Sereno, R. B. Tootell, and A. M. Dale, “High-resolution intersubject averaging and a coordinate system for the cortical surface,” Human Brain Mapping, vol. 8, no. 4, pp. 272–284, 1999.

[52] J. DiGuiseppi and P. Tadi, “Neuroanatomy, postcentral gyrus,” in StatPearls, StatPearls Publishing, 2025.

[53] K. Zilles and N. Palomero-Gallagher, “4.14 - the architecture of somatosensory cortex,” in The Senses: A Comprehensive Reference (Second Edition) (B. Fritzsch, ed.), pp. 225–260, Elsevier, 2020.

[54] T. Venot, Design and evaluation of a multimodal control of a robotic arm with a Brain Computer Interface. These de doctorat, Sorbonne université, Nov. 2023.

[55] R. M. El-Baba and M. P. Schury, “Neuroanatomy, frontal cortex,” in StatPearls, StatPearls Publishing, 2025.

[56] Y. Munakata, S. A. Herd, C. H. Chatham, B. E. Depue, M. T. Banich, and R. C. O’Reilly, “A unified framework for inhibitory control,” Trends in Cognitive Sciences, vol. 15, no. 10, pp. 453–459, 2011.

[57] K. Konstantopoulos and D. Giakoumettis, “Chapter 1 - basic knowledge on neuroanatomy and neurophysiology of the central nervous system,” in Neuroimaging in Neurogenic Communication Disorders (K. Konstantopoulos and D. Giakoumettis, eds.), pp. 1–30, Academic Press, 2023.

[58] F. Binkofski and G. Buccino, “Chapter 24 - the role of the parietal cortex in sensorimotor transformations and action coding,” in Handbook of Clinical Neurology (G. Vallar and H. B. Coslett, eds.), vol. 151 of The Parietal Lobe, pp. 467–479, Elsevier, 2018-01-01.

[59] J. Henderson, W. Choi, and S. Luke, “Morphology of primary visual cortex predicts individual differences in fixation duration during text reading,” Journal of Cognitive Neuroscience, vol. 26, pp. 2880–2888, 2014.

[60] E. T. Rolls, W. Cheng, and J. Feng, “The orbitofrontal cortex: reward, emotion and depression,” Brain Communications, vol. 2, no. 2, p. fcaa196, 2020-11-16.

[61] J. D. Wallis and M. F. S. Rushworth, “Chapter 22 - integrating benefits and costs in decision making,” in Neuroeconomics (Second Edition) (P. W. Glimcher and E. Fehr, eds.), pp. 411–433, Academic Press, 2014-01-01.

[62] E. Nozari, M. A. Bertolero, J. Stiso, L. Caciagli, E. J. Cornblath, X. He, A. S. Mahadevan, G. J. Pappas, and D. S. Bassett, “Macroscopic resting-state brain dynamics are best described by linear models,” Nature biomedical engineering, vol. 8, no. 1, pp. 68–84, 2024.

[63] F. Wendling, F. Bartolomei, J. J. Bellanger, and P. Chauvel, “Epileptic fast activity can be explained by a model of impaired gabaergic dendritic inhibition,” The European Journal of Neuroscience, vol. 15, no. 9, pp. 1499–1508, 2002.

[64] A. Puce and M. S. Hämäläinen, “A Review of Issues Related to Data Acquisition and Analysis in EEG/MEG Studies,” Brain Sci, vol. 7, May 2017.

[65] S. P. Ahlfors, J. Han, J. W. Belliveau, and M. S. Hämäläinen, “Sensitivity of MEG and EEG to Source Orientation,” Brain Topography, vol. 23, pp. 227–232, Sept. 2010.

[66] J. R. Wolpaw, D. J. McFarland, T. M. Vaughan, and G. Schalk, “The wadsworth center brain-computer interface (bci) research and development program,” IEEE Transactions on Neural Systems and Rehabilitation Engineering, vol. 11, no. 2, pp. 204–207, 2003.

[67] S. Taulu and J. Simola, “Spatiotemporal signal space separation method for rejecting nearby interference in meg measurements,” Physics in Medicine and Biology, vol. 51, no. 7, pp. 1759–1768, 2006.

[68] A. J. Bell and T. J. Sejnowski, “An information-maximization approach to blind separation and blind deconvolution,” Neural Comput, vol. 7, no. 6, pp. 1129–1159, 1995-11.

[69] R. Oostenveld, P. Fries, E. Maris, J.-M. Schoffelen, R. Oostenveld, P. Fries, E. Maris, and J.-M. Schoffelen, “FieldTrip: Open source software for advanced analysis of MEG, EEG, and invasive electrophysiological data,” Computational Intelligence and Neuroscience, vol. 2011, p. e156869, 2010-12.

[70] M. Fuchs, M. Wagner, and J. Kastner, “Boundary element method volume conductor models for EEG source reconstruction,” Clinical Neurophysiology, vol. 112, no. 8, pp. 1400–1407, 2001.

[71] A. Gramfort, T. Papadopoulo, E. Olivi, and M. Clerc, “OpenMEEG: opensource software for quasistatic bioelectromagnetics,” BioMedical Engineering OnLine, vol. 9, p. 45, 2010.

[72] M. Fuchs, M. Wagner, T. Köhler, and H. A. Wischmann, “Linear and nonlinear current density reconstructions,” Journal of Clinical Neurophysiology: Official Publication of the American Electroencephalographic Society, vol. 16, no. 3, pp. 267–295, 1999-05.

[73] F.-H. Lin, T. Witzel, S. P. Ahlfors, S. M. Stufflebeam, J. W. Belliveau, and M. S. Hämäläinen, “Assessing and improving the spatial accuracy in MEG source localization by depth-weighted minimum-norm estimates,” Neuroimage, vol. 31, no. 1, pp. 160–171, 2006-05.

[74] A. Gramfort, M. Luessi, E. Larson, D. A. Engemann, D. Strohmeier, C. Brodbeck, L. Parkkonen, and M. S. Hämäläinen, “MNE software for processing MEG and EEG data,” NeuroImage, vol. 86, pp. 446–460, 2014-02-01.

[75] F. Tadel, S. Baillet, J. Mosher, D. Pantazis, and R. Leahy, “Brainstorm: A user-firendly application for MEG/EEG analysis,” Computational Intelligence and Neuroscience, vol. 2011, 2011-01.

[76] J. C. Mazziotta, A. W. Toga, A. Evans, P. Fox, and J. Lancaster, “A probabilistic atlas of the human brain: theory and rationale for its development. the international consortium for brain mapping (ICBM),” NeuroImage, vol. 2, no. 2, pp. 89–101, 1995-06.

[77] R. S. Desikan, F. Ségonne, B. Fischl, B. T. Quinn, B. C. Dickerson, D. Blacker, R. L. Buckner, A. M. Dale, R. P. Maguire, B. T. Hyman, M. S. Albert, and R. J. Killiany, “An automated labeling system for subdividing the human cerebral cortex on MRI scans into gyral based regions of interest,” Neuroimage, vol. 31, no. 3, pp. 968–980, 2006-07.

[78] M.-C. Corsi, C. Gitton, J. Gonzalez-Astudillo, et al., “Understanding brain-computer interfaces training: a longitudinal and multimodal dataset,” Scientific Data, 2026.

[79] D. Wales and J. P. K. Doye, “Global optimization by basin-hopping and the lowest energy structures of lennard-jones clusters containing up to 110 atoms,” Journal of Physical Chemistry A, vol. 101, no. 28, pp. 5111–5116, 1997.

[80] P. Verma, S. Nagarajan, and A. Raj, “Stability and dynamics of a spectral graph model of brain oscillations,” Network Neuroscience, pp. 1–43, 07 2022.

[81] B. Rosner, R. J. Glynn, and M.-L. T. Lee, “The wilcoxon signed rank test for paired comparisons of clustered data,” Biometrics, vol. 62, no. 1, pp. 185–192, 2006.

[82] Y. Benjamini and D. Yekutieli, “The control of the false discovery rate in multiple testing under dependency,” The Annals of Statistics, vol. 29, no. 4, pp. 1165–1188, 2001.

[83] M. Friedman, “A comparison of alternative tests of significance for the problem of m rankings,” The annals of mathematical statistics, vol. 11, no. 1, pp. 86–92, 1940.

[84] J. C. Pinheiro and D. M. Bates, Mixed-Effects Models in S and S-PLUS. New York: Springer Science & Business Media, 2000.

