## Supplementary material for "Neural mechanisms of training in Brain-Computer Interface : A Biophysical modeling approach": Supplemenatry File

### Supplementary Document

### Results

**Replication with EEG** We replicated the condition effect analysis with EEG modeled data (refer to Fig. S4 and Table S6). No training effect has been observed (Friedman test, $p > 0.05$). Most of the significant conditions effects were observed from session 4. For $g_{\mathrm{ei}}$, such effects are involved in particular motor areas (left *precentral gyrus*), parietal lobe (left *inferior parietal* lobule) and occipital areas (bilateral *cuneus*). Similarly, for $g_{\mathrm{ii}}$, the bilateral cuneus showed significant effects. Lastly, for $\tau_{\mathrm{e}}$, the right supramarginal gyrus was significantly affected. These findings suggest session-specific variations in brain activity between MI and RS, mainly with session 4 that showed the most pronounced differences. Similarly to MEG, condition effects observed with $g_{\mathrm{ei}}$ and $g_{\mathrm{ii}}$ were associated with lower values in RS compared to MI, whereas $\tau_{\mathrm{e}}$ exhibited lower values in MI than in RS.

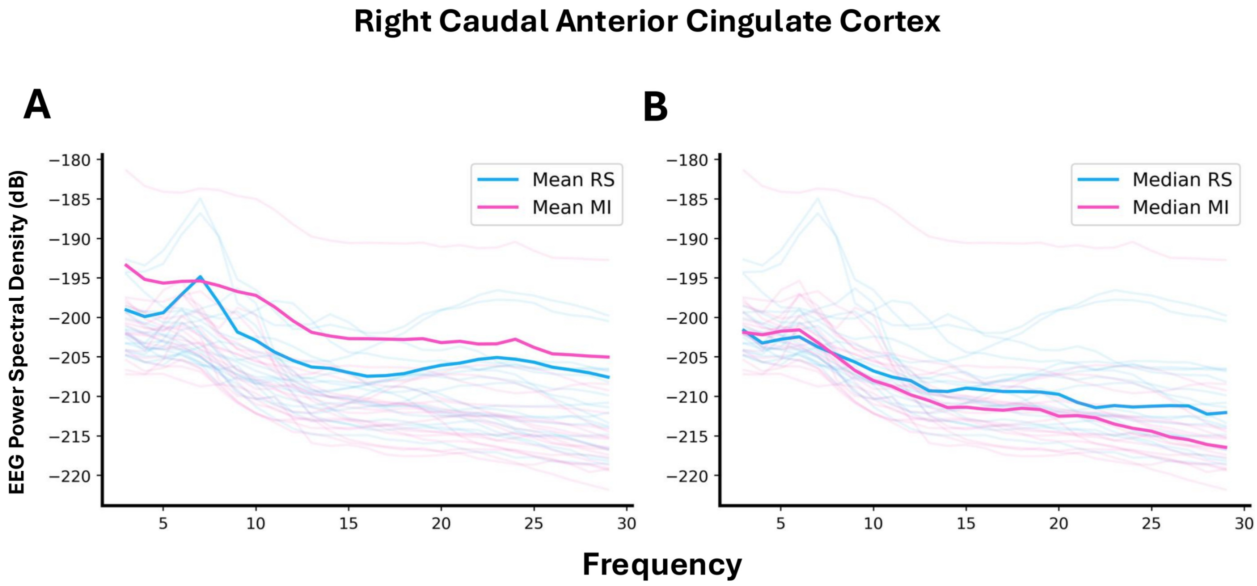


Figure S1: **Mean Vs. Median spectra:** Example comparison of median versus mean of spectra for the Right Caudal Anterior Cingulate region. Median spectra in B provides better representation of 19 individual spectra than mean spectra in A in both rest (RS) and motor-imagery (MI) condition, as few individual spectra skew the overall representation in case of mean.

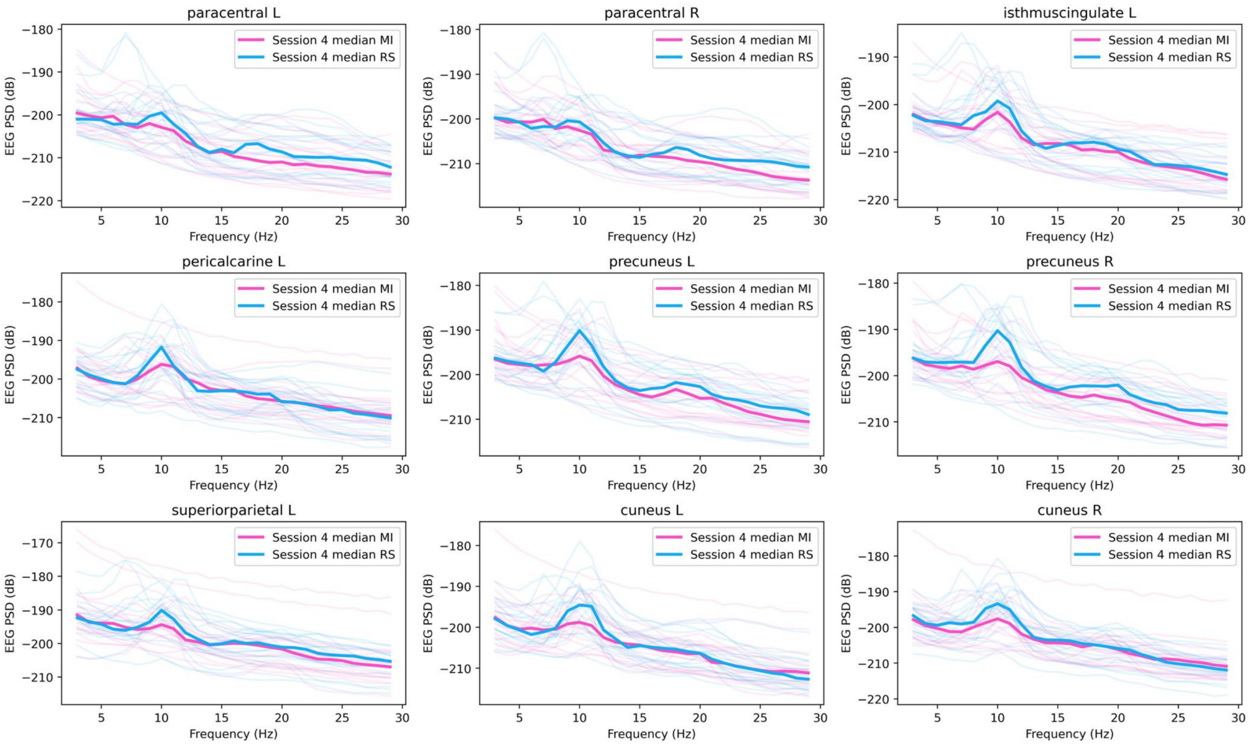


Figure S2: Representation of changes in power spectral density (PSD) in session 4 EEG along the frequency range (3-30 Hz) in a select few regions.

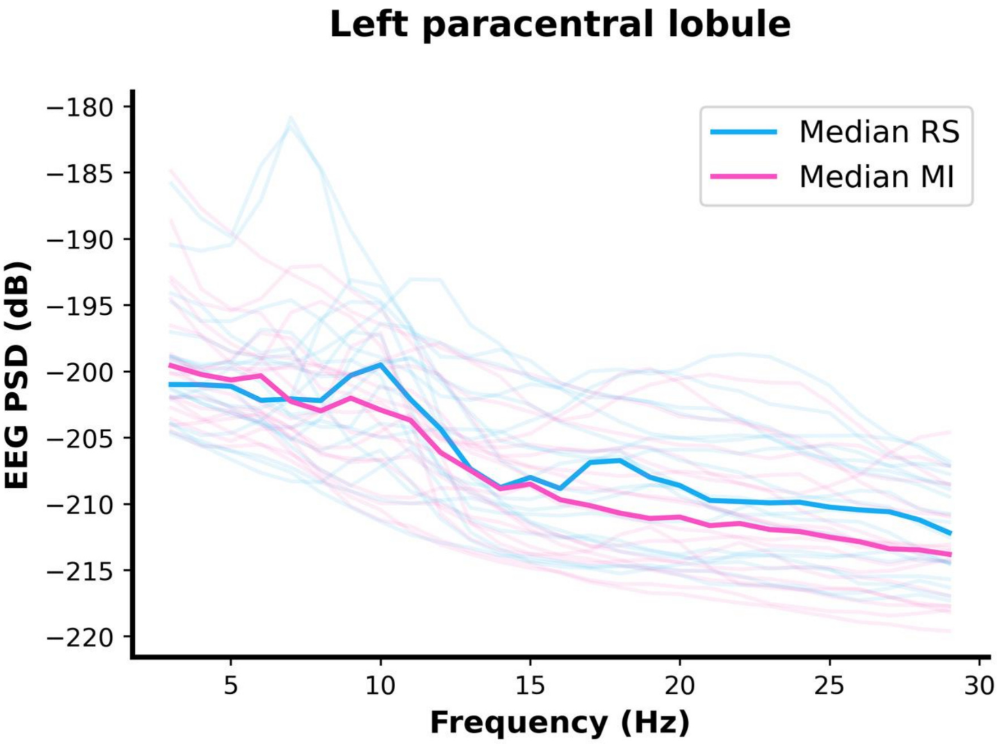


Figure S3: **Changes in EEG – Power Spectral Density (PSD) in Left paracentral lobule:** It shows that the changes in PSD in case of rest (RS) and motor-imagery (MI) condition, with median of the distribution of 19 spectra in both the conditions in bold. The alpha and beta peak can be observed in both the conditions.

Table S1: Sensitivity analysis of model parameters and input exponent for MEG modeled PSD. Spectra Sensitivity coefficient (mean ± standard error) across subjects, regions, and sessions.

| Perturbation ($\epsilon$) | $\tau_e$ | $\tau_i$ | $g_{ei}$ | $g_{ii}$ | $k$ |
|---|---|---|---|---|---|
| ***RS*** | | | | | |
| $\pm 5\%$ | $1.657 \pm 0.011$ | $0.838 \pm 0.014$ | $0.422 \pm 0.005$ | $0.356 \pm 0.008$ | – |
| $\pm 10\%$ | $1.581 \pm 0.010$ | $0.737 \pm 0.010$ | $0.398 \pm 0.005$ | $0.342 \pm 0.007$ | – |
| $\pm 20\%$ | $1.341 \pm 0.008$ | $0.647 \pm 0.007$ | $0.347 \pm 0.005$ | $0.305 \pm 0.006$ | – |
| $\pm 30\%$ | $1.108 \pm 0.006$ | $0.580 \pm 0.006$ | $0.282 \pm 0.004$ | $0.276 \pm 0.005$ | – |
| $\pm 40\%$ | $0.941 \pm 0.005$ | $0.511 \pm 0.005$ | $0.227 \pm 0.003$ | $0.250 \pm 0.004$ | – |
| ***MI*** | | | | | |
| $\pm 5\%$ | $1.364 \pm 0.010$ | $0.934 \pm 0.016$ | $0.352 \pm 0.005$ | $0.459 \pm 0.009$ | $0.165 \pm 0.002$ |
| $\pm 10\%$ | $1.295 \pm 0.009$ | $0.823 \pm 0.012$ | $0.337 \pm 0.005$ | $0.440 \pm 0.008$ | $0.165 \pm 0.002$ |
| $\pm 20\%$ | $1.042 \pm 0.006$ | $0.697 \pm 0.009$ | $0.285 \pm 0.004$ | $0.388 \pm 0.006$ | $0.167 \pm 0.002$ |
| $\pm 30\%$ | $0.811 \pm 0.005$ | $0.590 \pm 0.007$ | $0.232 \pm 0.003$ | $0.351 \pm 0.005$ | $0.169 \pm 0.002$ |
| $\pm 40\%$ | $0.639 \pm 0.004$ | $0.497 \pm 0.006$ | $0.192 \pm 0.003$ | $0.317 \pm 0.005$ | $0.173 \pm 0.003$ |

Table S2: Sensitivity analysis of model parameters and input exponent for EEG modeled PSD. Spectra Sensitivity coefficient (mean ± standard error) across subjects, regions, and sessions.

| Perturbation ($\epsilon$) | $\tau_e$ | $\tau_i$ | $g_{ei}$ | $g_{ii}$ | $k$ |
|---|---|---|---|---|---|
| ***RS*** | | | | | |
| $\pm 5\%$ | $1.927 \pm 0.013$ | $1.225 \pm 0.020$ | $0.389 \pm 0.006$ | $0.598 \pm 0.011$ | – |
| $\pm 10\%$ | $1.814 \pm 0.012$ | $1.029 \pm 0.015$ | $0.361 \pm 0.005$ | $0.567 \pm 0.010$ | – |
| $\pm 20\%$ | $1.488 \pm 0.009$ | $0.751 \pm 0.010$ | $0.307 \pm 0.005$ | $0.497 \pm 0.008$ | – |
| $\pm 30\%$ | $1.197 \pm 0.007$ | $0.637 \pm 0.007$ | $0.258 \pm 0.004$ | $0.432 \pm 0.007$ | – |
| $\pm 40\%$ | $0.994 \pm 0.005$ | $0.575 \pm 0.006$ | $0.216 \pm 0.003$ | $0.370 \pm 0.006$ | – |
| ***MI*** | | | | | |
| $\pm 5\%$ | $1.698 \pm 0.015$ | $1.394 \pm 0.024$ | $0.402 \pm 0.006$ | $0.722 \pm 0.014$ | $0.274 \pm 0.007$ |
| $\pm 10\%$ | $1.594 \pm 0.013$ | $1.163 \pm 0.017$ | $0.382 \pm 0.006$ | $0.687 \pm 0.013$ | $0.273 \pm 0.007$ |
| $\pm 20\%$ | $1.260 \pm 0.010$ | $0.856 \pm 0.010$ | $0.333 \pm 0.005$ | $0.597 \pm 0.010$ | $0.268 \pm 0.006$ |
| $\pm 30\%$ | $0.963 \pm 0.007$ | $0.691 \pm 0.008$ | $0.287 \pm 0.004$ | $0.511 \pm 0.008$ | $0.262 \pm 0.006$ |
| $\pm 40\%$ | $0.756 \pm 0.006$ | $0.595 \pm 0.006$ | $0.250 \pm 0.004$ | $0.438 \pm 0.007$ | $0.256 \pm 0.005$ |

Table S3: Sensitivity of MEG, MI-modeled spectra slope to model parameter and input exponent. Slope Sensitivity coefficient (mean ± standard error) across subjects, regions, and sessions.

| Perturbation ($\epsilon$) | $k$ | $\tau_e$ | $\tau_i$ | $g_{ei}$ | $g_{ii}$ |
|---|---|---|---|---|---|
| $\pm 10\%$ | $0.548 \pm 0.005$ | $0.525 \pm 0.005$ | $0.424 \pm 0.006$ | $0.235 \pm 0.004$ | $0.222 \pm 0.004$ |
| $\pm 20\%$ | $0.548 \pm 0.005$ | $0.434 \pm 0.004$ | $0.392 \pm 0.005$ | $0.193 \pm 0.003$ | $0.204 \pm 0.003$ |
| $\pm 30\%$ | $0.548 \pm 0.005$ | $0.374 \pm 0.004$ | $0.362 \pm 0.005$ | $0.151 \pm 0.003$ | $0.201 \pm 0.003$ |
| $\pm 40\%$ | $0.548 \pm 0.005$ | $0.348 \pm 0.003$ | $0.340 \pm 0.005$ | $0.120 \pm 0.002$ | $0.196 \pm 0.003$ |
| $\pm 50\%$ | $0.548 \pm 0.005$ | $0.322 \pm 0.003$ | $0.318 \pm 0.004$ | $0.098 \pm 0.002$ | $0.189 \pm 0.003$ |

Table S4: Sensitivity of EEG, MI-modeled spectra slope to model parameter and input exponent. Slope Sensitivity coefficient (mean ± standard error) across subjects, regions, and sessions.

| Perturbation ($\epsilon$) | $k$ | $\tau_e$ | $\tau_i$ | $g_{ei}$ | $g_{ii}$ |
|---|---|---|---|---|---|
| $\pm 10\%$ | $1.859 \pm 0.397$ | $1.562 \pm 0.147$ | $1.314 \pm 0.160$ | $0.696 \pm 0.147$ | $1.434 \pm 0.462$ |
| $\pm 20\%$ | $1.859 \pm 0.397$ | $0.941 \pm 0.079$ | $0.950 \pm 0.075$ | $0.486 \pm 0.078$ | $0.929 \pm 0.139$ |
| $\pm 30\%$ | $1.859 \pm 0.397$ | $0.939 \pm 0.094$ | $0.836 \pm 0.080$ | $0.407 \pm 0.084$ | $0.878 \pm 0.172$ |
| $\pm 40\%$ | $1.859 \pm 0.397$ | $0.788 \pm 0.075$ | $0.583 \pm 0.031$ | $0.355 \pm 0.074$ | $0.769 \pm 0.128$ |
| $\pm 50\%$ | $1.859 \pm 0.397$ | $0.806 \pm 0.094$ | $0.559 \pm 0.054$ | $0.308 \pm 0.063$ | $0.706 \pm 0.123$ |

Table S5: Significant ROIs ($P_{\text{FDR}} < 0.05$) across MEG sessions, grouped by lobe

| **Session** | $g_{\text{ei}}$ | $g_{\text{ii}}$ | $\tau_{\text{e}}$ |
|---|---|---|---|
| Session 1 | Parietal (Right inferior and superior Parietal), Occipital (Left lateral Occipital), Frontal (Left pars Opercularis) | Temporal (Left transverse Temporal) | Temporal (Left banks of superior temporal sulcus (STS), Left Fusiform, Left inferior and middle Temporal, Left pars Opercularis, Left and right transverse Temporal), Occipital (Left and right Cuneus, Left and right Pericalcarine), Frontal (Left and right caudal middle Frontal, Left and right Precentral, Right superior Frontal), Parietal (Left Paracentral, Left and right Postcentral, Right Precuneus, Left and right Supramarginal), Limbic (Right Entorhinal, Right posterior Cingulate) |
| Session 2 | Nil | Temporal (Left transverse Temporal) | Occipital (Left and right Cuneus, Left and right Pericalcarine), Temporal (Left Fusiform, Left Insula, Left pars Opercularis, Right pars Triangularis, Left transverse Temporal, Right transverse Temporal), Frontal (Right medial Orbitofrontal, Right Precentral), Parietal (Left Paracentral, Right Postcentral, Right Precuneus, Left superior Parietal, Right Supramarginal), Limbic (Left posterior Cingulate, Left rostral anterior Cingulate) |
| Session 3 | Nil | Parietal (Left inferior parietal, Left lateral Occipital), Frontal (Left rostral middle frontal), Temporal (Left and right transverse temporal), Limbic (Right posterior Cingulate) | Temporal (Left banks STS, Left Inferior temporal, Left pars Opercularis, Left and right transverse temporal), Occipital (Left and right Cuneus, Left pericalcarine), Frontal (Right caudal middle Frontal, Left and right Precentral, Left superior Frontal), Parietal (Left inferior Parietal, Left and right Postcentral, Left and right superior Parietal, Right superior temporal, Left and right Supramarginal), Limbic (Right posterior Cingulate) |
| Session 4 | Nil | Parietal (Right postcentral), Temporal (Left transverse temporal) | Temporal (Left banks STS, Left transverse temporal), Occipital (Left cuneus, Left pericalcarine), Frontal (Right precentral), Parietal (Left and right inferior parietal, Left and right paracentral, Left and right postcentral, Left and right supramarginal), Limbic (Left entorhinal, Right isthmus cingulate, Left and right posterior cingulate, Left and right precuneus) |

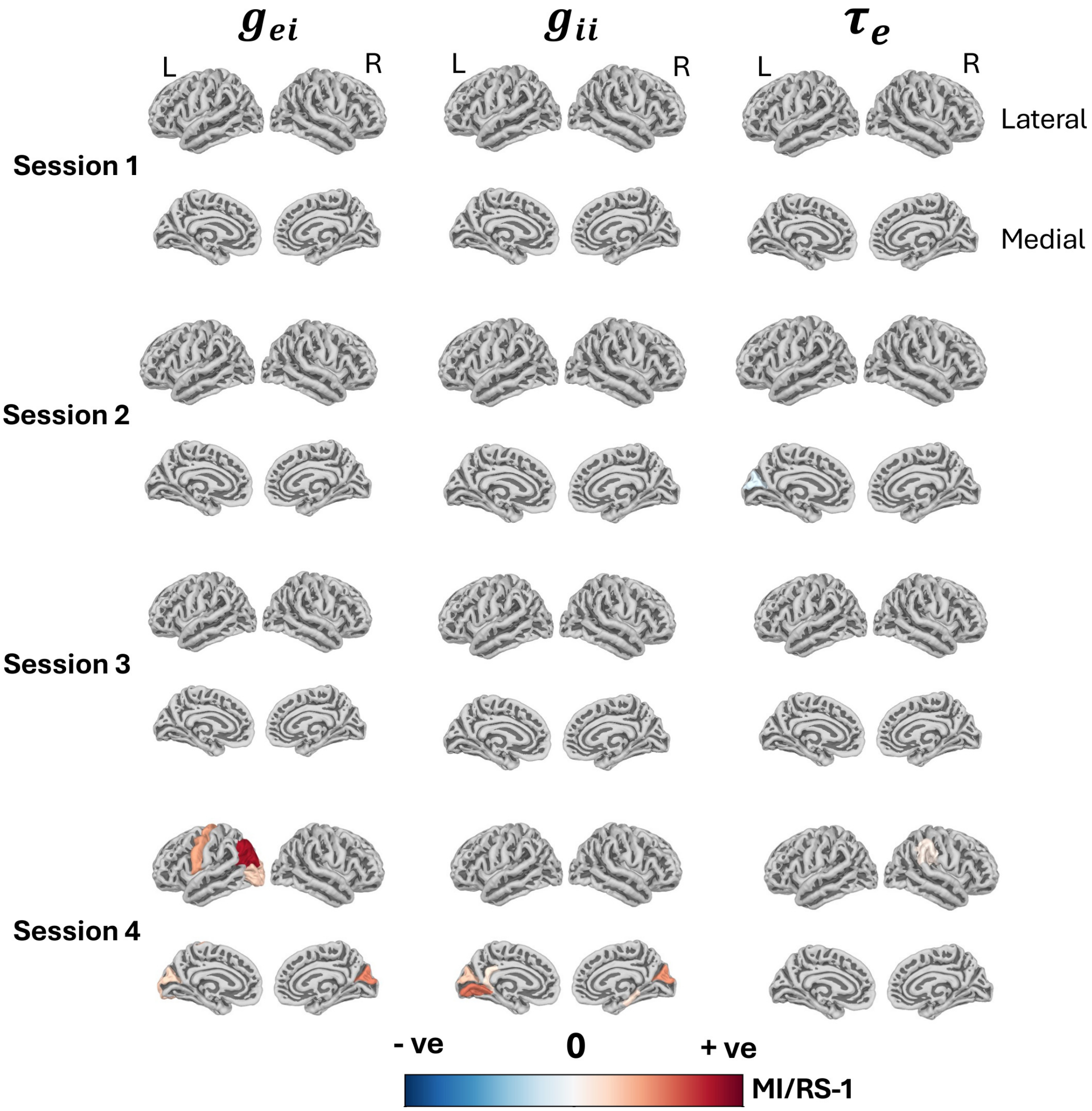


Figure S4: **EEG – based condition effect analysis in BCI learning:** The highlighted colored regions show statistically significant ($p_{\mathrm{FDR}} < 0.05$) condition effect for MI Vs RS for the three EEG fitted model parameters ($g_{\mathrm{ei}}$, $g_{\mathrm{ii}}$, $\tau_{\mathrm{e}}$) respectively in every session. This shows functional reorganization across sessions as various cortical regions are getting activated for the MI BCI task in the 19 subjects' group. Minimum and maximum range of mean (MI/RS-1) for the parameters across sessions are (-0.38, +184.74), (-0.418, +374.87), (-0.365, +1.449) for $g_{\mathrm{ei}}$, $g_{\mathrm{ii}}$, $\tau_{\mathrm{e}}$ respectively.

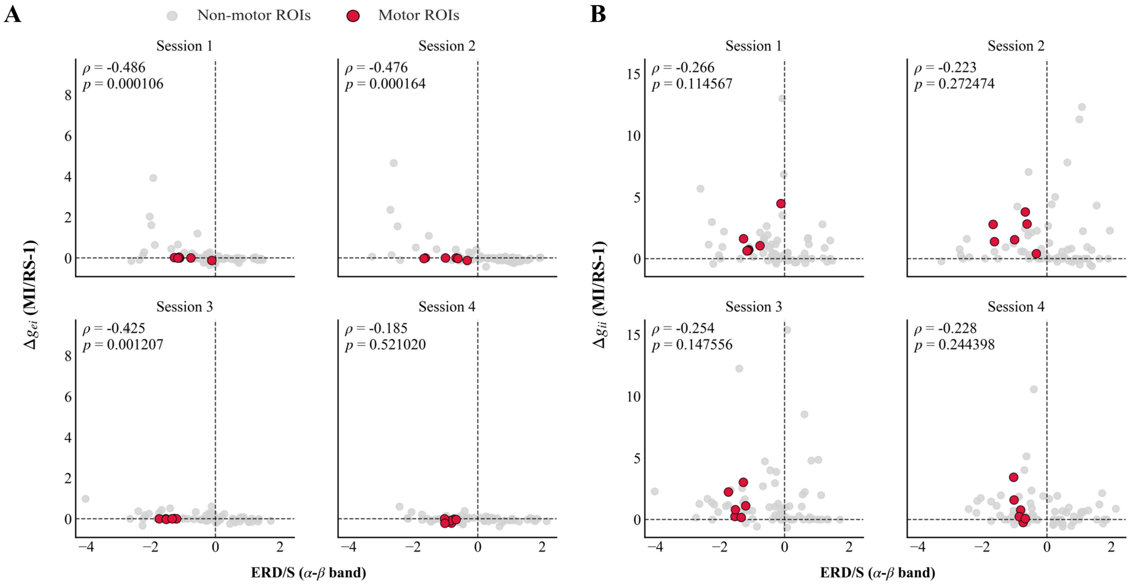


Figure S5: **Association between ERD/S and neural gains at the global level.** Plot A shows significant negative association between ERD/S and $g_{\mathrm{ei}}$ from session 1 to session 3 (Bonferroni corrected $p < 0.005$) and lost significance by session 4 (Bonferroni corrected $p > 0.05$). Plot B shows no significant association between ERD/S and $g_{\mathrm{ii}}$ for any sessions (Bonferroni corrected $p > 0.05$).

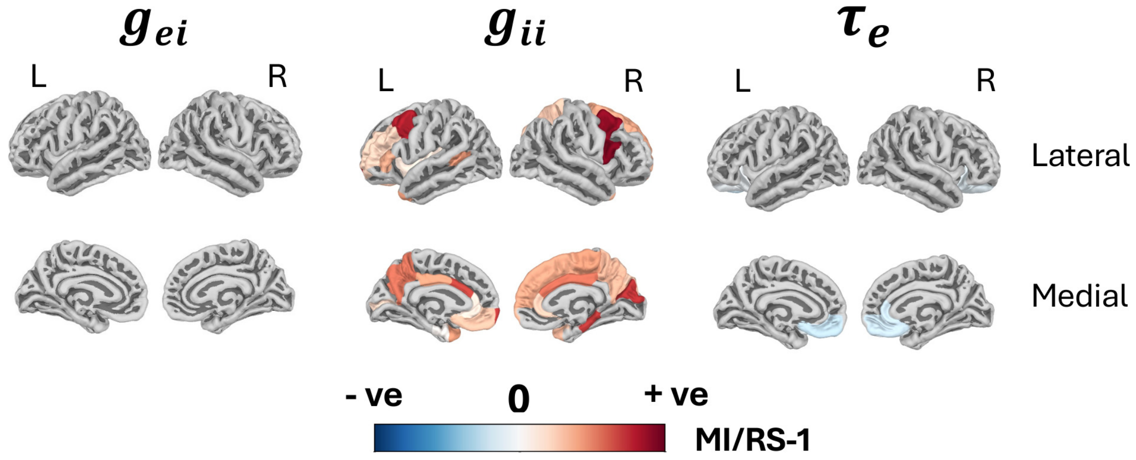


Figure S6: **Condition effect analysis for the independent dataset:**The highlighted colored regions show statistically significant ($p_{\mathrm{FDR}} < 0.05$) condition effect between MI and RS for the three EEG fitted model parameters ($g_{\mathrm{ei}}$, $g_{\mathrm{ii}}$, $\tau_{\mathrm{e}}$) respectively. This show how various cortical regions are getting activated for MI BCI task. Minimum and maximum range of mean (MI/RS-1) for the parameters across sessions are (-0.287, +81.57), (-9.275, +416.3), (-0.36, +1.448) for ($g_{\mathrm{ei}}$, $g_{\mathrm{ii}}$, $\tau_{\mathrm{e}}$) respectively.

Table S6: Significant ROIs ($P_{\text{FDR}} < 0.05$) across EEG sessions

| **Session** | $g_{\text{ei}}$ | $g_{\text{ii}}$ | $\tau_{\text{e}}$ |
|---|---|---|---|
| Session 1 | Nil | Nil | Nil |
| Session 2 | Nil | Nil | Left Cuneus |
| Session 3 | Nil | Nil | Nil |
| Session 4 | Left and right Cuneus, Left inferior Parietal, Left lateral Occipital, Left Precentral | Left and right Cuneus, Left isthmus Cingulate, Left Lingual, Right Parahippocampal | Right Supramarginal |
